# Knowledge-guided Bayesian optimization using pre-trained LLMs speeds up the identification of superior genotypes from germplasm collection

**DOI:** 10.64898/2026.06.28.735149

**Authors:** Kosuke Hamazaki, Koji Tsuda

## Abstract

**Background:** Germplasm collections contain wide genetic diversity that is valuable for plant breeding, but conducting phenotypic evaluation for all genotypes in field trials is rarely feasible. Bayesian optimization offers a way to decide, season by season, which genotypes to cultivate in order to identify superior genotypes with fewer evaluations. However, standard Bayesian optimization commonly starts from randomly selected genotypes and mainly relies on surrogate models built from marker genotype information, while the text-based passport information that accompanies germplasm is not fully used. We examined whether pre-trained large language models can provide prior knowledge that improves these decisions in germplasm evaluation.

**Results:** We constructed a large-language-model-guided Bayesian optimization framework that introduces large language models into two parts of the Bayesian optimization workflow. In zero-shot warmstarting, a large language model proposes initial genotypes using passport information such as cultivar name, country of origin, and subpopulation, optionally together with principal component scores derived from genome-wide single-nucleotide-polymorphism markers. In addition, we evaluated a large-language-model-based surrogate model that predicts phenotypic values for untested genotypes using in-context learning from previously evaluated genotypes. Using a rice germplasm panel and two target traits (seed number per panicle for maximization and protein content for minimization), we compared strategies. For seed number per panicle, zero-shot warmstarting with a general-purpose instruction-following model reduced the number of evaluated genotypes needed to reach the best genotype, whereas improvements were small for protein content. When genomic information was available, Gaussianprocess-based Bayesian optimization was the strongest overall approach, while the large-language-model-based surrogate model outperformed random baselines and was competitive in some settings. When genomic information was not available, predictions based on passport information improved efficiency compared with fully random strategies.

**Conclusions:** Pre-trained large language models can inject useful agronomic knowledge into Bayesian optimization for germplasm evaluation, particularly by improving early-stage genotype selection, and can also support optimization when genomic information is unavailable. As models better handle long genomic sequences together with passport information, large-language-model-guided Bayesian optimization may become a practical and explainable decision-support approach for agricultural optimization.

## Introduction

Plant breeding increasingly requires the development of cultivars tolerant to drought, heat, salinity, and other stresses to cope with future climate change and uncertainty in growing environments [1, 2]. Achieving this requires moving beyond the variation present within existing breeding populations and actively considering the introgression of external genetic resources, such as wild relatives and landraces with high diversity [3–5]. In particular, germplasm collections encompass diverse genotypes, including wild relatives, and are therefore expected to be valuable for breeding [6]. Indeed, germplasm has been collected for many crop species and is increasingly organized into databases as curated collections; in some cases, such as rice [7, 8] and soybean [9, 10], genome sequences for individual genotypes within these resources have become publicly available. However, the vast size of these collections means that phenotypic evaluation remains insufficient in many crop species, creating a major bottleneck to their effective use in breeding.

To make effective use of germplasm in breeding, it is essential to characterize the potential of each genotype through phenotypic evaluation in field trials and related experiments [11]. However, because the number of candidate genotypes is often extremely large, evaluating all of them in the field at once is rarely feasible. A more practical approach is to cultivate and evaluate a subset of genotypes each year, gradually accumulating phenotypic information [12]. When expert knowledge about the genotypes is unavailable, however, evaluations may proceed in a random order, which is highly inefficient in terms of the number of trials required. To address this problem, Tanaka and Iwata proposed an annual evaluation strategy based on Bayesian optimization (BO) to efficiently identify superior genotypes from large germplasm collections [13, 14]. BO can iteratively propose the next candidates to evaluate by balancing “exploitation,” which focuses on promising regions based on existing information, and “exploration,” which targets uncertain regions to gain new information, thereby enabling optimization with relatively few trials [15, 16]. Accordingly, BO has also been applied in plant-breeding contexts beyond germplasm evaluation, including the optimization of breeding strategies [17–19]. Nevertheless, conventional BO still leaves room for improvement; for example, the initial genotypes to be cultivated are often chosen at random [20].

Further improvements in the efficiency of such optimization methods will likely benefit from incorporating rapidly advancing AI technologies. In particular, large language models (LLMs) have attracted substantial attention because they learn patterns of knowledge and linguistic regularities from massive text corpora, enabling context-aware text generation and reasoning [21, 22]. Incorporating LLMs into predictive and decision-making frameworks makes it possible to handle not only numerical data but also natural-language information [22–24]. This is especially appealing for agricultural datasets, which often contain rich text-based information such as passport data describing country of origin, subpopulation structure, and collection-site narratives. Accordingly, diverse applications of LLMs in agriculture are emerging [25]. In particular, research on domain-specific LLMs trained intensively on agricultural knowledge has expanded, including PLLaMa [26], AgriGPT [27], and evaluations of GPT-4 on agricultural exam-style questions [28]. Related work in the broader plant-breeding context includes SeedBench, which measures knowledge and reasoning in breeding and seed science [29], and PlantGPT, which integrates phenotypic and gene-function data with retrieval to improve response quality as a “virtual expert” [30]. These efforts reflect ongoing progress in building knowledge-intensive and evaluation infrastructures for breeding [31].

LLMs are also expected to improve the efficiency of optimization methods that rely on predictive models [32, 33]. For instance, LLAMBO [34] integrates LLMs into the BO framework; by using LLMs to select initial candidates and as surrogate models to estimate unobserved points, it has been shown to increase the efficiency of hyper-parameter optimization in machine learning [34–36]. This LLM-guided BO paradigm is gradually being extended to multiple domains, including design, materials discovery, and chemistry, suggesting that combining knowledge representations encoded by language models with optimization algorithms may be a promising way to accelerate exploration [37–39].

Although research on LLMs in agriculture has begun and the usefulness of LLM-guided BO has been demonstrated in several fields, concrete applications of LLM-guided BO to specific agricultural problems remain limited. In particular, at key stages such as initial candidate selection, effectively eliciting the domain knowledge encoded in LLMs may provide substantial benefits in agriculture, where background knowledge (e.g., crop biology, cultivation regions, management conditions, and variety information) strongly influences decision-making. Therefore, in this study, we use germplasm evaluation as a case study and introduce LLMs into two components of BO: (i) selecting genotypes to be cultivated initially and (ii) developing surrogate models to predict genotypic values for untested genotypes using historical field-trial data. Through these implementations, we aim to improve optimization efficiency and examine the potential for applying LLMs to agricultural optimization problems.

## Materials and Methods

In this study, we used R version 4.5.2 for data formatting and visualization [40], and specifically used the R ggplot2 package version 4.0.0 for visualization [41].

### Bayesian optimization for identifying superior genotypes

#### Problem formulation

In this study, following the settings in Tanaka and Iwata, we consider a scenario in which we aim to identify superior genotypes from a germplasm collection with known marker genotypes [13, 14]. Identifying superior genotypes requires phenotypic evaluation through field trials, which is time-consuming and costly. In particular, since the number of genotypes that can be evaluated per season is limited, it is necessary to efficiently identify superior genotypes with a limited number of evaluations.

We formulate this germplasm evaluation problem as an optimization problem (Fig. 1). The search space is a high-dimensional genotype space **W**∈ **{**0, 1, 2} ^*N*×*M*^ represented by genome-wide SNP marker information. We want to solve a discrete optimization problem of selecting *L*^(*t*)^ ∈ N genotypes from *N* ∈ N genotypes in the germplasm panel for field trial evaluation each year to identify the best genotype as quickly as possible. The objective function is represented by the phenotypic value of each genotype **y** = *f* (**W**) ∈ R^*N*^, and we aim to identify the genotype with the maximum or minimum phenotypic value depending on the breeding objective.

**Fig. 1.**
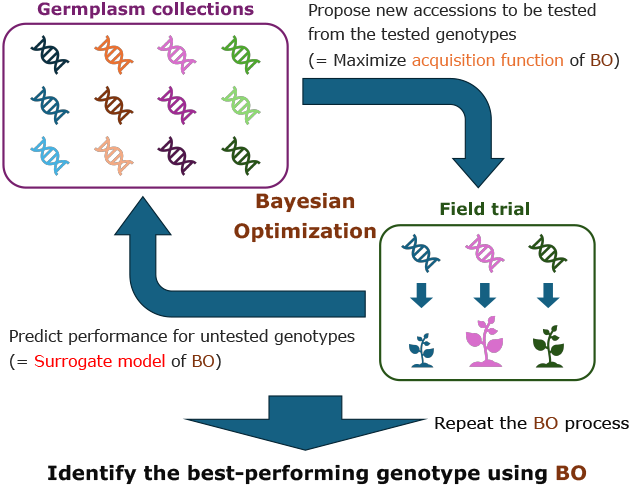
Conceptual illustration of germplasm evaluation and the application of Bayesian optimization.

This problem presents two main challenges. First, the objective function *f* is unknown. In other words, the exact relationship between marker genotypes and phenotypes is unclear, making this a black-box optimization problem where gradient-based methods cannot be applied. Second, evaluating the objective function *f*, that is, conducting phenotypic evaluation, is highly costly. Therefore, minimizing the number of function evaluations through an appropriate black-box optimization method is essential.

#### Bayesian optimization framework

BO is a method for efficiently performing global optimization of black-box functions with high evaluation costs [15, 16]. It is well-suited to the problem we address in this study: efficiently identifying superior genotypes from germplasm collections. BO searches for optimal solutions by repeating a cycle of four steps: selecting initial observation points (zero-shot warmstarting), evaluating proposed points, constructing and updating a surrogate model, and selecting the next evaluation points by maximizing an acquisition function. Below, we explain how BO applies to identifying superior genotypes in germplasm collections [13, 14] (Fig. 2).

**Fig. 2.**
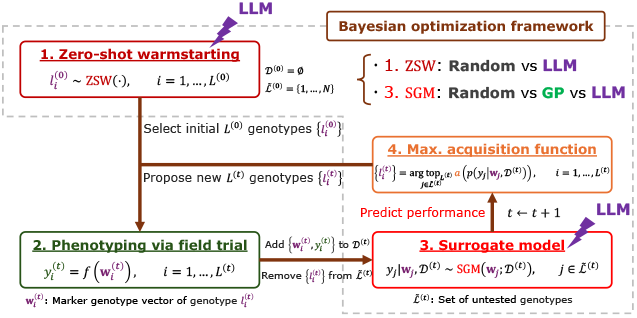
Conceptual comparison of conventional Bayesian optimization and LLM-guided Bayesian optimization for germplasm evaluation. The figure illustrates the Bayesian optimization (BO) framework using germplasm evaluation as an example. In this study, BO consists of four steps: 1. selecting genotypes to be evaluated in the initial field trial (zero-shot warmstarting), 2. phenotyping through field trials, 3. predicting the genetic ability of untested genotypes using a surrogate model, and 4. selecting the next genotypes to evaluate in subsequent field trials by maximizing an acquisition function. We focus on incorporating LLMs into steps 1 and 3 and compare the resulting approaches with conventional BO.

1. Zero-shot warmstarting (Fig. 2-1) First, we initialize the dataset *D*^(*t*)^ containing marker genotype and phenotype information for tested genotypes, and the set 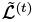 containing untested genotype IDs as follows:

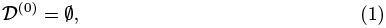

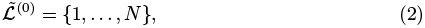

where ∅ is the empty set and *N* ∈ N is the number of genotypes. Next, we select *L*^(0)^ ∈ N genotypes to be initially tested in field trials, which are necessary for constructing the surrogate model of BO, as in the following equation.

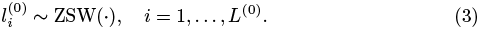

Here, 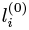 is *i*-th ID of the selected genotypes, and ZSW is a function that samples genotype IDs for initial field trials from the set 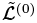 of untested genotype IDs. In standard BO, ZSW function is often replaced with random selection.
2. Phenotyping via field trial (Fig. 2-2) Next, we consider conducting field trials on the selected genotypes 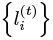 to obtain phenotype data. The phenotype 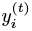 corresponding to genotype ID 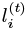 is genetically controlled, so it can be expressed as a function of the corresponding marker genotype 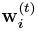:

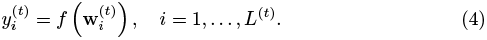

The challenge is that the function *f* representing the relationship between phenotypes and marker genotypes is unknown and stochastic rather than deterministic. BO addresses this by iteratively approximating *f* with a surrogate model and maximizing an acquisition function. In this study, rather than conducting actual field trials, we performed simulations where all known open data were initially masked and hidden, with only phenotype data for the selected genotypes becoming available through field trials. When phenotypes 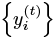 are obtained from field trials, we update the dataset *D*^(*t*)^ of tested genotypes by adding the current field trial results 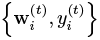. We also update the set 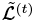 of untested genotype IDs by removing 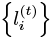.
3. Surrogate model (Fig. 2-3) Next, we predict the phenotypic value *y*_*j*_ of an untested genotype 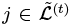 using the *n*^(*t*)^ data points tested so far, D^(*t*)^ = {**W**^(*t*)^, **y**^(*t*)^}, as follows:

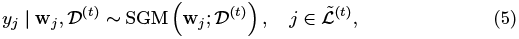

where SGM is a function that returns a probability distribution from the tested data *D*^(*t*)^ and the input marker genotype **w**_*j*_ using a surrogate model. One of the most popular surrogate models is the Gaussian process (GP), which constructs a prediction model in the following normal distribution [42].

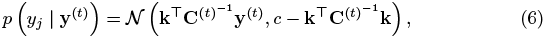

Where 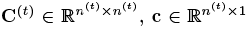, and *c* ∈ R are obtained by partitioning the variance-covariance matrix 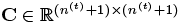 of 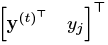 as follows:

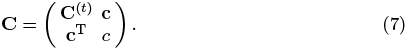

This covariance matrix **C** is expressed as follows:

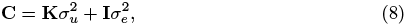

where 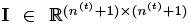 is the identity matrix, and 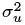 and 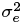 are the genetic variance and error variance estimated from the data, respectively.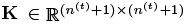 is a Gaussian kernel calculated from the marker genotypes 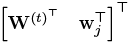, and the (*a, b*) component of **K** is calculated as follows:

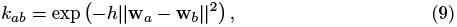

where *h* is a hyperparameter that determines the bandwidth. In this study, we used the value calculated from the median of the off-diagonal components of the distance matrix between marker genotypes. In this way, GP use marker genotype information to calculate the predictive distribution of phenotypic values for untested genotypes. In this study, the implementation of GP was performed using Python’s gpytorch package version 1.14.1 [43].
4. Maximize acquisition function (Fig. 2-4) Once the predictive distribution of phenotypic values is calculated, we compute the acquisition function for each untested genotype. The acquisition function is calculated by considering the balance between the mean and variance predicted by the Gaussian process. To identify the genotype with the maximum phenotypic value, we select the genotype that maximizes the acquisition function as the next candidate for field trials. That is, after updating *t* ← *t* + 1, we select the next *L*^(*t*)^ ∈ N genotypes for field trials in descending order of their acquisition function values:

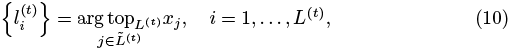

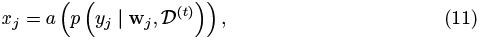

where 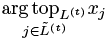 is a function that returns the IDs of the top *L*^(*t*)^ genotypes from the untested set 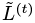 with the largest *x*_*j*_ values, and *a* is a function that computes the acquisition function from the predictive distribution calculated by the Gaussian process. There are several candidates for specific acquisition functions, including UCB (Upper Confidence Bound), PI (Probability of Improvement), and EI (Expected Improvement). In this study, we adopted UCB [44], which is calculated from the mean *µ*_*j*_ and variance 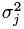 of the predictive distribution as follows:

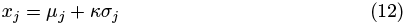

where *κ* is a hyperparameter that controls the balance between exploration and exploitation. In this study, we set *κ* = 2. Note that when we want to identify the genotype with the minimum phenotypic value, we use

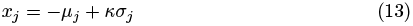

to select the top *L*^(*t*)^ genotypes *j* with the largest *x*_*j*_ values as 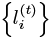.
5. Repeat until convergence By repeating steps 2–4 until convergence, BO efficiently identifies the genotype with the maximum or minimum phenotypic value. In this study, since the maximum or minimum value in the population is known, we repeated steps 2–4 above at least until BO reached this value.

### Idea of LLM-guided BO

As proposed by Liu et al., the standard Bayesian optimization described above has room for improvement [34–36]. In this study, we therefore focused on the following two key points when identifying superior individuals from germplasm collections and attempted to integrate LLMs into BO (Fig. 2). LLMs have already learned extensive knowledge about agriculture and genetics, and BO that leverages this prior knowledge in agronomy may be able to identify superior genotypes more efficiently.

First, while conventional BO often randomly selects initial points in zero-shot warmstarting, there is room to leverage prior domain knowledge in agronomy for this selection. Specifically, by providing the LLM with passport information such as variety names, country of origin, and subpopulation information along with the target trait to be maximized or minimized, we can connect this information to the LLM’s prior knowledge to propose promising initial candidate points for BO (Fig. 3). This enables exploration of promising regions from the start of optimization and is expected to reduce the number of evaluations needed to discover the best genotype.

**Fig. 3.**
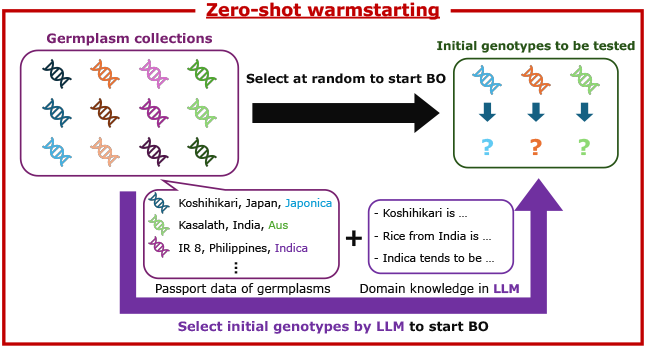
Conceptual illustration of the rationale for incorporating LLMs into zero-shot warmstarting in Bayesian optimization for germplasm evaluation.

Second, we examine an approach that uses LLMs as surrogate models (Figure S1). In this study, we train the LLM through in-context learning on observed data from field trials, allowing it to predict phenotypic values for untested genotypes. Since LLMs can handle not only genomic information but also string representations such as passport information through natural language, they may be more effective than conventional Gaussian processes in complex germplasm evaluation problems. However, as discussed in the Discussion section, technical challenges exist, including prompt length limitations and methods for representing genomic data.

In this subsection, we explained the idea of LLM-guided BO for evaluating germplasm collections. More detailed implementation of the LLM-guided BO is described in the Implementation and evaluation of LLM-guided BO subsection.

## Materials and problem settings

### Germplasm collection and Marker genotype data

In this study, we used a germplasm collection of 413 rice (Oryza sativa L.) accessions from the “Rice diversity” project [45]. This collection was collected from 85 countries, such as the United States, India, and China, and comprises six subpopulations: tropical japonica, temperate japonica, indica, aus, aromatic, and admixture (others). We downloaded the 44K SNP set by Zhao et al. [45] as marker genotypes from the link “44K SNP set” at http://www.ricediversity.org/data/index.cfm. We then extracted only biallelic SNPs with minor allele frequencies ≥0.05 and missing rates ≤0.1 from the downloaded data using PLINK 1.9 [46, 47] and VCFtools version 0.1.13 [48], and then imputed them using Beagle version 4.1 [49, 50]. This process yielded 29,556 SNPs, which were used for subsequent analyses. This marker genotype data is identical to that used in Hamazaki and Iwata [17].

### Phenotype data

For phenotype data, we used both real and simulated data to evaluate the influence of the LLM’s domain knowledge on the target trait.

#### Real data

For real data, we used seed number per panicle and protein content from the 34 traits published in Zhao et al. [45]. Seed number per panicle is a trait to be maximized, with data available for 376 accessions. In contrast, protein content is a trait to be minimized, with data available for 393 accessions.

#### Simulated data

For the above two traits, the relevant information may be included in an LLM’s domain knowledge. In contrast, it is an interesting question how the performance of LLM-guided BO changes when the LLM lacks domain knowledge of the target trait. Therefore, in this study, we also considered a setting in which the LLM has no such domain knowledge and simulated phenotypes accordingly.

For the simulated traits, we generated phenotypic values under three scenarios using the following model:

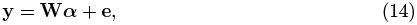

where 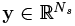 is a vector of simulated phenotypic values, 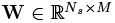 is a matrix of genotype scores for QTNs, ***α*** ∈ R^*M*^ is a vector of QTN effects, 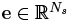 is an error vector, and *N*_*s*_ = 413. We sampled QTN effects ***α*** from a univariate normal distribution. We then determined the error variance from the ratio to the genetic variance based on the given heritability *h*^2^ and sampled errors from a normal distribution.

The three scenarios differed in the number of QTNs and heritability as follows. Scenario 1: *M* = 5, 000, *h*^2^ = 0.7, Scenario 2: *M* = 5, 000, *h*^2^ = 0.3, and Scenario 3: *M* = 50, *h*^2^ = 0.7. Due to computational constraints, we simulated phenotype values only once for each scenario and performed BO with maximization as the objective in all cases.

### Simulation settings

Using the rice data described above, we identified genotypes with maximum or minimum phenotypic values for each trait through BO, following the procedure in the Bayesian optimization for identifying superior genotypes subsection. Here, the total number of genotypes *N* varied by target trait: *N* = 376 for seed number per panicle, *N* = 393 for protein content, and *N* = 413 for simulated traits. In this study, we fixed the number of initially selected genotypes and the number evaluated in field trials at each iteration to *L*^(0)^ = *L*^(*t*)^ = 10. As mentioned in the Bayesian optimization framework subsection, we did not conduct actual field trials for the selected genotypes. Instead, we ran simulations where only the phenotypic information for selected genotypes became accessible from the masked open data. We repeated BO 100 times for each trait and evaluated each strategy based on changes in the maximum or minimum phenotypic values of tested genotypes. Details of the compared strategies are in the Strategies compared in the study subsection.

### Implementation and evaluation of LLM-guided BO

As described in the Idea of LLM-guided BO subsection, we introduced LLMs into both zero-shot warmstarting and the surrogate model. This subsection describes the implementation details of the LLMs.

### Six LLMs used in the study

In this study, we used the following six models available from Hugging Face (https://huggingface.co/): PLLaMa-7b-instruct [26] (PLLaMa), CropSeek-LLM [51] (CropSeek), gemma-7b-it [52] (Gemma), Meta-Llama-3-8B-Instruct [53] (Llama), Mistral-7B-Instruct-v0.1 [54] (Mistral), and openchat-3.5-1210 [55] (Openchat). PLLaMa is based on LLaMa-2 and trained on over 1.5 million plant science-related academic papers, making it rich in knowledge of plant and agricultural science [26]. CropSeek is fine-tuned on agriculture-related text based on DeepSeek-R1-14B [56] and contains information on crop descriptions, diseases, and agricultural techniques [51]. Thus, while these two models are domain-specific models that specialize in agricultural-related information, the remaining four models are foundation models that excel at instruction-following. We downloaded each model locally from Hugging Face and executed it in a local environment. For running these LLMs, we used Python’s torch package version 2.6.0 [57], transformers package version 4.56.2 [58], and sentence transformers package version 5.1.1 [59].

When using these LLMs, prompt length is a constraint, making it difficult to directly input long sequences like genomic data. To address this, we performed principal component analysis on marker genotype information consisting of 29,556 SNPs. We then used the principal component scores from the first to fifth components as representative genomic information for each genotype as input to the LLMs. Principal component analysis was performed using version 1.7.2 of Python’s scikit-learn package [60].

### Zero-shot Warmstarting (ZSW)

In zero-shot warmstarting (ZSW), we select *L*^(0)^ = 10 genotypes from *N* genotypes for initial field trials. When using an LLM for this selection, we must provide passport and genomic information for each genotype. However, including all this information creates excessively long prompts that prevent proper inference. To address this, we first used Retrieval-Augmented Generation (RAG) [24] to select the top 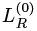 texts semantically similar to “genotypes suitable for initializing BO” (query) from the large amount of information available for each genotype. We then divided the information on these 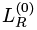 genotypes into chunks and had the LLM select 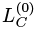 genotypes from each chunk. The number of chunks was automatically determined based on token length constraints and ranged from approximately 10 to 20. Finally, from the genotypes selected in each chunk, we had the LLM select *L*^(0)^ = 10 genotypes through one more round of instruction. The parameters 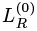 and 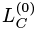 were set to 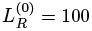 and 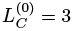 when using genomic information, and to 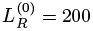 and 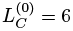 when not using it.

The input prompts to the LLM consisted of the following elements:

- **Task description**: Explanation of using BO to identify superior genotypes from *N* genotypes in germplasm collections, with instruction to select *L*^(0)^ = 10 genotypes suitable for initializing BO.
- **Trait information**: Name of the target trait and breeding objective (maximize/minimize).
- **Passport information**: Line or cultivar name and subpopulation information for each genotype. Country of origin was also included when genomic information was not used.
- **Genomic information**: Principal component scores from the first to fifth components obtained from principal component analysis of marker genotypes.
- **Output format**: Instruction to return the indices of candidate genotypes in Python list format.

For ZSW, we tested all six models described in the Six LLMs used in the study subsection.

### Surrogate model (SGM)

The surrogate model (SGM) predicts phenotypic values *y*_*j*_ of untested genotypes 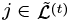 based on previous field trial results *D* ^(t)^= {**W**^(t)^, **y** ^(t)^**}**. When using an LLM for predictions, we employed in-context learning by providing *D*^(*t*)^ as examples within the prompt rather than fine-tuning with *D*^(*t*)^. However, as the number of tested genotypes *n*^(*t*)^ increases, the information required for examples grows, causing prompt length limitations. To address this, when 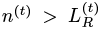, we used RAG to screen *D*^(*t*)^ and select the top 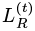 data points semantically similar to “efficient candidate points for performing predictions with SGM.” Predictions were then made using min 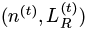 data points, with 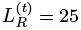.

The specific input prompts to the LLM consisted of the following elements:

- **Task description**: Explanation to predict the phenotype of the target trait from subpopulation information and genomic information (or subpopulation and country of origin information when genomic data is unavailable) using in-context learning based on examples of previous trial results.
- **Trait information**:Name of the target trait and breeding objective (maximize/minimize).
- **Observed data**: List of cultivar names, subpopulation information, genomic information (or subpopulation and country of origin information when genomic data is unavailable), and corresponding phenotypic values for genotypes evaluated so far.
- **Prediction target**: Cultivar name, subpopulation information, and genomic information (or subpopulation and country of origin information when genomic data is unavailable) for a single genotype whose phenotypic value is to be predicted.
- **Output format**: Instruction to provide the predicted phenotypic value as a single numerical value.

Here, predictions by the SGM were performed 100 times individually for each genotype. Since LLMs are probabilistic generative models, these 100 responses differ each time, allowing them to be regarded as a pseudo-heuristic predictive distribution. In this study, based on the predictive distribution calculated from the 100 responses, we computed the acquisition function introduced in the Bayesian optimization framework subsection and selected the genotypes for field trials in the next cycle.

Due to computational time constraints, we used only two LLMs in the SGM: Llama and Mistral.

### Strategies compared in the study

To evaluate the performance of LLM-guided BO, we prepared seven strategies (LLG, LL, LG, LRG, LR, RG, RR) as shown in Table 1. These strategies differ in their use of Random or LLM for ZSW, their use of Random, GP, or LLM for SGM, and whether they utilize genomic information. When GP is used for SGM, genomic information is required, so there is no strategy using GP without genomic information. The RR strategy uses Random for both ZSW and SGM and does not require genomic information, so “Use Genome” is automatically set to No. Additionally, when LLMs are used for both ZSW and SGM (LLG, LL), the same model was used for both steps.

**Table 1.**
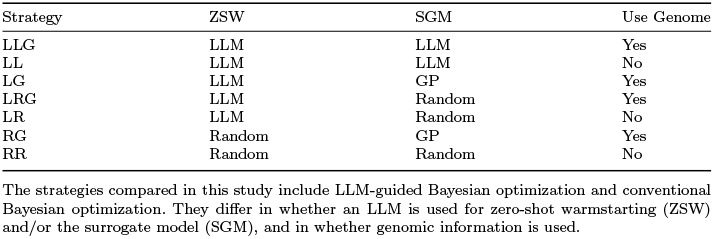
Strategies compared in this study.

| Strategy | ZSW | SGM | Use Genome |
| --- | --- | --- | --- |
| LLG | LLM | LLM | Yes |
| LL | LLM | LLM | No |
| LG | LLM | GP | Yes |
| LRG | LLM | Random | Yes |
| LR | LLM | Random | No |
| RG | Random | GP | Yes |
| RR | Random | Random | No |

The strategies compared in this study include LLM-guided Bayesian optimization and conventional Bayesian optimization. They differ in whether an LLM is used for zero-shot warmstarting (ZSW) and/or the surrogate model (SGM), and in whether genomic information is used.

Now, in the subsequent Result section, to clearly visualize the impacts of introducing LLMs into ZSW and SGM, we compared strategies separately rather than all at once. To evaluate the impact of LLMs in ZSW, we compared LG (six LLMs) and RG. To evaluate the impact of LLMs in SGM with genomic information, we compared three strategies: LLG (Llama or Mistral), LG, and LRG. To evaluate the impact of LLMs in SGM without genomic information, we compared three strategies: LL (Llama), LR, and RG.

Each strategy was evaluated primarily by the number of field trials required to identify the best genotype. We also evaluated prediction accuracy for unknown genotypes at each step using the correlation coefficient between observed and predicted values.

## Results

### Results for real data

#### Comparison of zero-shot warmstarting strategies

First, we focused on ZSW and evaluated the impact of introducing LLMs into ZSW. As described in the Strategies compared in the study, we assessed the RG and six LLM-based LGs in Table 1 based on the number of field trials required to identify the best genotype (Fig. 4). The x-axis represents the number of genotypes evaluated in field trials, and the y-axis shows the maximum (for seed number per panicle) or minimum (for protein content) phenotypic value among the genotypes evaluated up to that point. Thus, a strategy is considered better when its curve is closer to the upper left (for seed number per panicle) or the lower left (for protein content), indicating faster identification of the best genotype.

**Fig. 4.**
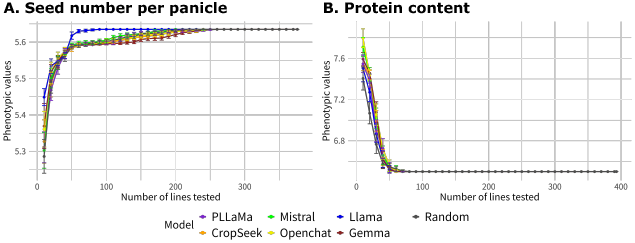
Comparison of LLMs for zero-shot warmstarting (ZSW): improvement in the maximum/minimum phenotypic value among selected genotypes for (A) seed number per panicle and (B) protein content. This figure evaluates how quickly each model can identify the best genotype by plotting the change in the maximum/minimum phenotypic value among selected genotypes (y-axis) against the number of genotypes evaluated in field trials so far (x-axis). Therefore, curves located closer to the upper left indicate superior performance. We compared ZSW using LLMs, two agriculture-oriented models and four general-purpose models, with random selection. Model abbreviations and corresponding line colors are as follows: PLLaMa (purple), CropSeek (orange), Mistral (light green), Openchat (yellow), Llama (blue), Gemma (brown), and Random (black).

From the seed number per panicle results, we found that the LG using Llama as the LLM identified the best genotype in the population the fastest (Fig. 4A). In contrast, LGs using other LLMs performed similarly to RG. Specifically, the average number of genotypes evaluated until identifying the best genotype was more than 120 for models other than Llama, whereas it was 53.2 for Llama, indicating that Llama required nearly 2.5 times fewer genotype evaluations than RG, which required 131.8 evaluations on average (Supplementary Table 1). Among the non-Llama LGs, only Mistral identified the best genotype earlier than RG on average; the other four models, including the agriculture-specific LLM, performed worse than RG. We also examined how SGM accuracy changed over time and found that, particularly in the early to mid stages (up to when approximately 180 genotypes had been cultivated), the LG using Llama as the LLM showed higher accuracy than the other models (Supplementary Fig. 2A).

To investigate why the LG using Llama as the LLM achieved superior performance for seed number per panicle, we repeated the LLM-based selection of genotypes to be initially tested in field trials 100 times and counted how often each genotype was selected across the 100 runs. We visualized these counts on a principal component plot based on marker genotypes (Fig. 5, Supplementary Fig. 3). In this plot, point color indicates the rice subpopulation, and point size reflects the selection frequency across the 100 runs; larger points indicate genotypes that were more frequently chosen as initial candidates for cultivation. Points selected at least once are shown as filled circles, whereas those never selected are shown as open circles. Under RG (Random), because each genotype is chosen at random, every genotype is selected on average about 100 ×*L*^(0)^*/N* = 100 ×10*/*376 ≈2.66 times, resulting in points of similar size (Fig. 5C). In contrast, under PLLaMa, point sizes varied, indicating that some genotypes were selected much more frequently than others; a similar tendency was observed for all LGs (Fig. 5AB, Supplementary Fig. 3). Notably, Llama showed the largest variation in point size, and some genotypes were selected as initial candidates more than 50 times out of 100 runs (Fig. 5A).

**Fig. 5.**
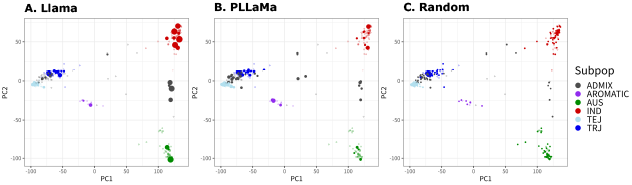
Genotype selection frequency in three zero-shot warmstarting models for seed number per panicle. Selection frequency across 100 LLM-guided ZSW runs for the initial field trials, shown on a principal component (PC) plot derived from marker genotypes (PC1 vs PC2). Here, we compared three models: (A) Llama, (B) PLLaMa, and (C) Random. Point color denotes rice subpopulation, and point size is proportional to the number of times each genotype was selected (0–100). Filled circles indicate genotypes selected at least once; open circles indicate genotypes never selected.

For the minimization of protein content, however, all LGs and RG showed nearly equivalent performance (Fig. 4A). Specifically, the average number of genotypes evaluated until identifying the best genotype was lowest for Llama (37.2), while even Openchat, which required the most evaluations, needed only 42.2; therefore, no substantial differences were observed among models (Supplementary Table 2). In the corresponding principal component plots, as in Fig. 5, models differed somewhat in the degree of variation in point sizes; however, for Llama in particular, the variation was smaller than in the seed number per panicle case (Supplementary Fig. 4). In terms of SGM accuracy, as in the seed number per panicle case, the LG using Llama as the LLM showed slightly higher accuracy than the other models, particularly in the early to mid stages (up to approximately 180 cultivated genotypes) (Supplementary Fig. 2B).

#### Comparison of surrogate models

Next, to evaluate the impact of introducing LLMs into SGM with genomic information, we compared LLGs using Llama or Mistral as the LLM with LG and LRG in terms of convergence speed, following the same procedure as in Fig. 4 (Fig. 6, Supplementary Fig. 5). When using Llama, LG identified the best genotype the fastest for both traits, followed by LLG and then LRG (Fig. 6). In other words, although LLG that incorporated an LLM as SGM outperformed LRG, which selects the next candidates at random, it did not outperform LG, which uses the conventional Gaussian process. This trend was also supported by the number of genotypes evaluated until the best genotype was identified (Supplementary Table 3 for seed number panicle and Supplementary Table 4 for protein content). Moreover, for both traits, SGM accuracy was higher with Gaussian process–based predictions than with LLM-based predictions (Supplementary Fig. 6). When using Mistral, protein content showed a similar trend to Llama, whereas for seed number per panicle, LLG identified the best genotype slightly earlier than LG. Thus, depending on the LLM used, SGM may be able to outperform conventional methods (Supplementary Fig. 5, Supplementary Table 5 for seed number per panicle and Supplementary Table 6 for protein content).

**Fig. 6.**
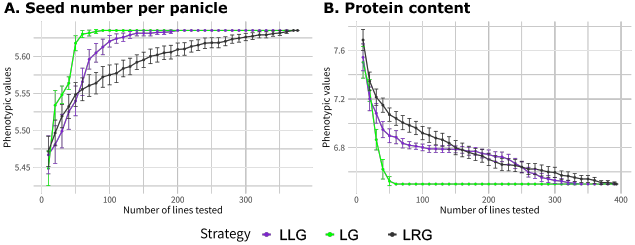
Comparison of the Llama-based surrogate model (SGM) and the conventional SGM when genomic information is available: improvement in the maximum/minimum phenotypic value among selected genotypes for (A) seed number per panicle and (B) protein content. As in Fig. 4, the x-axis denotes the number of genotypes already evaluated in field trials, and the y-axis denotes the maximum/minimum phenotypic value among the genotypes evaluated so far. Here, we compared three strategies using genomic information: LLG (purple), which uses an LLM (Llama) for both ZSW and SGM; LG (light green), which uses Llama for ZSW and a Gaussian process for SGM; and LRG (black), which uses Llama for ZSW and Random for SGM.

When an LLM is used as the SGM, it can make predictions using only passport information, enabling BO without relying on genomic data. Therefore, to evaluate the impact of introducing an LLM into SGM when genomic information is unavailable, we compared LL (using Llama as the LLM) with LR and RR, both of which randomly select the next genotype to cultivate (Fig. 7). Note that when genomic information is not used, GP-based prediction cannot be performed; thus, comparison with strategies that adopt GP as the SGM is not possible. As a result, for both traits, performance was best in the order LL, LR, and RR (Fig. 6). On average, LL succeeded in finding the best genotype 1.24 times faster than LR for seed number per panicle (Supplementary Table 7) and 1.21 times faster for protein content (Supplementary Table 8). These findings suggest that, even in situations where genomic information cannot be obtained due to cost or other constraints, LLM-based prediction using passport information may contribute to improving the efficiency of BO.

**Fig. 7.**
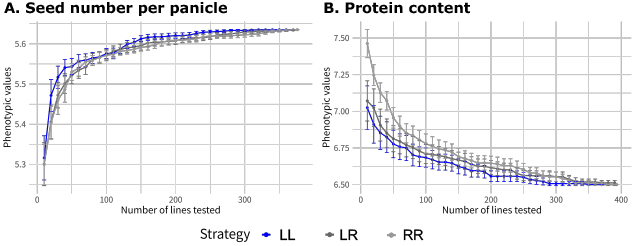
Comparison of the Llama-based surrogate model (SGM) and the conventional SGM when genomic information is unavailable: improvement in the maximum/minimum phenotypic value among selected genotypes for (A) seed number per panicle and (B) protein content. As in Fig. 4, the x-axis denotes the number of genotypes already evaluated in field trials, and the y-axis denotes the maximum/minimum phenotypic value among the genotypes evaluated so far. Here, we compared three strategies using genomic information: LL (blue), which uses an LLM (Llama) for both ZSW and SGM; LR (black), which uses Llama for ZSW and Random for SGM; and RR (gray), which uses Random for both ZSW and SGM.

### Comparison of zero-shot warmstarting strategies for simulated data

We next examined in more detail the incorporation of LLMs into ZSW, which was particularly effective when seed number per panicle was the target trait. Using the simulated phenotypic data generated in the Simulated data subsection, we compared RG and six LLM-based LGs in Table 1 using the same procedure as in the Comparison of zero-shot warmstarting strategies subsection (Figs 8–10).

**Fig. 8.**
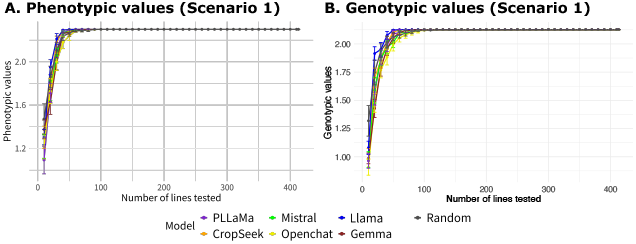
Comparison of LLMs for zero-shot warmstarting (ZSW): improvement in the maximum phenotypic/genotypic value among selected genotypes for the trait simulated under Scenario 1. As in Fig. 4, the x-axis represents the number of genotypes that have already been evaluated in field trials, and the y-axis represents the maximum (A) phenotypic value or (B) genotypic value among the genotypes evaluated so far. Model abbreviations and the line colors corresponding to each model are the same as in Fig. 4.

In scenario 1, which assumes a high-heritability trait governed by many QTLs, the LG using Llama performed slightly better than LGs using other LLMs and RG under both phenotypic value–based and genotypic value-based evaluations; however, the differences were small (Fig. 8). In contrast, in scenario 2, which assumes a low-heritability trait governed by many genes, the LG using Llama substantially outperformed all other models under both evaluation criteria (Fig. 9); notably, this trend was evident from the first field trial. In scenario 3, which assumes a high-heritability trait governed by a small number of genes, as in scenario 1, the LG using Llama slightly outperformed the other models, but no major differences were observed among models (Fig. 10). Under phenotypic value-based evaluation, all strategies showed an apparent plateau around 2.25, followed by further improvement after the number of cultivated genotypes reached 150 (Fig. 10A). By contrast, under genotypic value-based evaluation, the best genotype had already been identified by the time 100 genotypes had been cultivated (Fig. 10B). This discrepancy arises because the genotype with the maximum phenotypic value does not necessarily coincide with the genotype with the maximum genotypic value; therefore, from a breeding perspective, strategies that enable earlier identification of the genotype with the highest genotypic value are preferable.

**Fig. 9.**
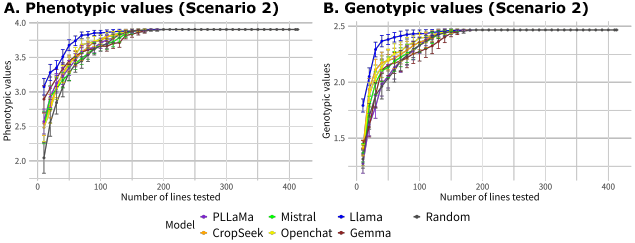
Comparison of LLMs for zero-shot warmstarting (ZSW): improvement in the maximum phenotypic/genotypic value among selected genotypes for the trait simulated under Scenario 2. As in Fig. 4, the x-axis represents the number of genotypes that have already been evaluated in field trials, and the y-axis represents the maximum (A) phenotypic value or (B) genotypic value among the genotypes evaluated so far. Model abbreviations and the line colors corresponding to each model are the same as in Fig. 4.

**Fig. 10.**
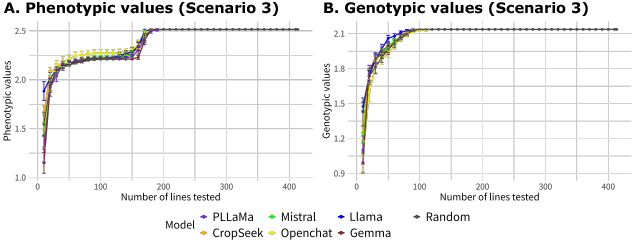
Comparison of LLMs for zero-shot warmstarting (ZSW): improvement in the maximum phenotypic/genotypic value among selected genotypes for the trait simulated under Scenario 3. As in Fig. 4, the x-axis represents the number of genotypes that have already been evaluated in field trials, and the y-axis represents the maximum (A) phenotypic value or (B) genotypic value among the genotypes evaluated so far. Model abbreviations and the line colors corresponding to each model are the same as in Fig. 4.

We also generated principal component plots for each trait under scenarios 1 to 3, analogous to Fig. 5 (Supplementary Figs 8 - 10). The results indicated no substantial differences in the genotypes selected by each model across scenarios. Compared with other models, Llama, which showed particularly strong performance, tended to repeatedly select a smaller subset of genotypes. These frequently selected genotypes were drawn broadly from all subpopulations, with a particular emphasis on IND and ADMIXED.

## Discussion

### Interpretations of the results

In this study, to explore the applicability of LLMs in agriculture, we investigated how LLMs could be incorporated into a BO framework for agricultural problems. Specifically, we introduced (i) LLM-based zero-shot warmstarting (ZSW) initialization and (ii) an LLM-based surrogate model (SGM) into BO, and evaluated their effectiveness using a germplasm evaluation problem as an example.

First, in the ZSW results for seed number per panicle, Llama achieved the best performance (Fig. 4A). The number of times each genotype was selected showed the greatest variability for Llama, indicating that Llama repeatedly selected certain genotypes many times (Fig. 5A). By contrast, in the ZSW results for protein content, where Llama performed comparably to the other models, we did not observe a tendency for a subset of genotypes to be repeatedly selected (Supplementary Fig. 4A). These observations suggest that Llama may retain some domain knowledge related to seed number per panicle and, based on that knowledge, may select the same genotypes repeatedly with a certain level of confidence.

Across these ZSW results, the agriculture-oriented LLMs (PLLaMa, CropSeek) did not outperform the other LLMs (Fig. 4A). Because these models were fine-tuned on a large amount of agriculture-related literature [26, 51], this outcome may seem counterintuitive at first glance. However, training on a large corpus of agricultural literature does not necessarily provide knowledge relevant to seed number per panicle or protein content, and model accuracy may therefore not improve [61]. Even more importantly, we confirmed cases in which these agriculture-oriented LLMs exhibited poorer instruction-following performance than general-purpose LLMs and thus could not adequately respond to concrete exploration instructions, such as “propose promising initial genotype names for performing BO efficiently.” Continual fine-tuning of LLMs has been reported to cause catastrophic forgetting, in which previously acquired knowledge and abilities for reasoning and reading comprehension are degraded [62]. In addition, instruction tuning and alignment based on human feedback have been shown to be important for obtaining responses that follow user instructions [61, 63]. Therefore, to effectively leverage agriculture-oriented LLMs for BO, it is necessary to consider fine-tuning strategies that are designed not only to acquire agricultural knowledge but also to maintain instruction-following performance.

Next, we discuss the ZSW results for simulated traits. Because LLMs are not expected to possess domain knowledge about simulated traits, the impact of introducing LLMs into ZSW is likely to be smaller than for real traits. Nevertheless, particularly in Scenario 2, which assumes low-heritability traits controlled by many genes, using Llama contributed substantially to improving BO efficiency (Fig. 9). In contrast, in Scenarios 1 and 3, which assume high-heritability traits, although Llama slightly outperformed the other models, we did not observe large differences (Figs 8 and 10). This can be explained by the fact that when heritability is high, the SGM is sufficiently accurate that any model can quickly identify the best genotype, whereas when heritability is low, the choice of initial genotypes more strongly influences SGM accuracy.

The number of times each LLM selected particular genotypes as candidates for initial field trials differed little across scenarios (Supplementary Figs 8 - 10). This is reasonable because, for simulated traits, the trait name carries little information, leaving the models with little basis for distinguishing among traits. Notably, however, compared with the other models, Llama tended to proactively and repeatedly select representative genotypes across subpopulations. This suggests that Llama selected not only highly promising genotypes but also a diverse set of genotypes. In scenarios such as Scenario 2, which assumes traits controlled by many QTLs, constructing an SGM from diverse genotypes may facilitate predictions that capture population structure, thereby improving SGM accuracy from an early stage. Overall, these results indicate that even when an LLM does not retain domain knowledge about the target trait, selecting a sufficiently diverse set of initial genotypes may, depending on the trait, contribute to improved BO efficiency.

We next discuss the results obtained when an LLM was introduced as the SGM. When Llama was used as the SGM, performance was worse than when a GP was used as the SGM for both seed number per panicle and protein content (Fig. 6). Although prediction accuracy was close to zero (Supplementary Fig. 6), Llama still outperformed the LRG baseline, which selects the next genotype to cultivate at random. This suggests that using an LLM to incorporate genomic and passport information can still be beneficial. One possible explanation for the improved BO efficiency despite low predictive accuracy is that, even if the predicted values themselves are inaccurate, the model can still estimate predictive variance with reasonable accuracy based on which genotypes have already been tested, which can be useful when computing the acquisition function.

A likely reason for the low accuracy in this study is that the LLM could not accept original genomic information as an extremely long sequence as input; instead, we represented it using principal component scores. Conventional GP models can fully exploit genomic information for prediction and may therefore have an advantage that more than compensates for the inability to use passport information. This limitation may be mitigated as genome language models (gLMs), which have recently attracted attention in areas such as gene function prediction [64, 65], continue to develop and as predictive models that can handle both long sequences and passport information emerge [66]. Such advances could improve the predictive accuracy of LLM-based SGMs and potentially allow them to outperform GPs. Indeed, we already observed cases in which LLMs achieved performance exceeding that of GPs (e.g., when Mistral was used as the SGM for seed number per panicle; Supplementary Fig. 5), suggesting that the potential of using LLMs will continue to increase depending on the model.

Finally, we discuss the results when genomic information is unavailable. Because conventional GP-based approaches use only genomic information as explanatory variables, they cannot run BO with a GP as the SGM when genomic data are unavailable. In contrast, LLMs can use passport information as input, enabling BO with an LLM-based SGM even without genomic data. When we applied an LLM using only passport information to ZSW and SGM, BO performance improved slightly relative to random selection (Supplementary Fig. fig:7), suggesting that passport information contains some signal that is useful for efficiently identifying the best genotype. However, given an improvement on the order shown in Supplementary Fig. fig:7, one might argue that breeding experts’ judgment could be more efficient. Nonetheless, because not everyone necessarily has expert-level knowledge about the genotypes of interest, the ability for anyone to make informed decisions as long as passport information is available is a major advantage of our proposed approach. Moreover, although we used only passport information in this study, incorporating additional genotype-related information (e.g., pedigree information) could further improve the accuracy of LLM-based SGMs and potentially achieve performance comparable to, or even exceeding, that of experts.

### Advantages and future perspectives of LLM-guided BO

As discussed in the previous subsection, the framework we propose for incorporating LLMs into BO has the advantage that it can be used by anyone, provided that the necessary information is available. This is because an LLM can partially substitute for the agricultural domain knowledge typically held by experts and can use that knowledge to guide the search. In addition, unlike conventional BO, the LLM can be prompted to explain why it selected particular genotypes, including the initial genotypes to cultivate and the next genotype to evaluate. In the future, this could enable an interactive workflow in which BO is performed through dialogue with an LLM, much like asking an expert to justify their choices. Such explainable BO has attracted attention not only in agriculture but also in many other fields [36, 67, 68]. Beyond explainability, the proposed approach is also promising in terms of performance. As more agricultural data accumulate and as LLMs continue to advance, this framework is expected to contribute increasingly to improving BO efficiency.

Finally, we consider other potential applications of the proposed LLM-guided BO framework in agriculture. This framework is not limited to germplasm evaluation. It can be applied more broadly to settings in which the goal is to identify optimal candidates under a limited number of trials. Because agriculture is a domain where prior knowledge, such as knowledge of target traits and environmental responses, plays an important role, LLMs may be able to guide early-stage exploration by using such knowledge as cues. For example, this approach could support BO in decision-making tasks such as selecting crossing parents and designing breeding programs. It could also support cultivation management decisions related to fertilization, irrigation, and sowing timing by integrating information that an LLM can utilize, such as historical data, field conditions, characteristics of varieties, and empirical rules. In summary, LLM-guided BO can be extended to a wide range of decision-making and optimization problems in agriculture, and its usefulness is expected to grow further as data and knowledge continue to accumulate.

## Conclusion

In this study, we addressed the challenge of identifying superior genotypes from a germplasm collection with a limited number of phenotypic evaluations. To this end, we proposed and validated a framework that integrates the prior domain knowledge of LLMs into BO. Specifically, by introducing LLMs into both (i) ZSW, which proposes genotypes for initial field evaluation, and (ii) SGM, which predicts phenotypes for unevaluated individuals, we demonstrated the potential to incorporate knowledge into decision-making from the early stages of the search.

The main conclusion is that incorporating the prior domain knowledge of LLMs into BO can improve the efficiency of germplasm evaluation (i.e., identifying the best individual with fewer evaluations). This approach is not limited to germplasm evaluation; it can be extended to a broad range of agronomic optimization problems in which domain knowledge critically affects outcomes, such as breeding program design, selection of mating pairs, and decision-making regarding crop-management. Looking ahead, as models capable of jointly handling long and diverse information, such as genome sequences (e.g., gLMs) and pedigree data, continue to advance, and as agronomic data accumulation progresses, LLM-guided BO is expected to have a major impact on agriculture as a highly accurate and practically useful decision-support foundation.

## Supplementary information

**Supplementary Fig. 1. Conceptual illustration of the rationale for incorporating LLMs into surrogate models in Bayesian optimization for germplasm evaluation**.

**Supplementary Fig. 2. Comparison of LLMs for zero-shot warmstarting (ZSW): change in the surrogate model accuracy for (A) seed number per panicle and (B) protein content**. The figure plots the change in the surrogate model accuracy (y-axis) against the number of genotypes evaluated in field trials so far (x-axis). Model abbreviations and the line colors corresponding to each model are the same as in Fig. 4.

**Supplementary Fig. 3. Genotype selection frequency in four zero-shot warmstarting models for seed number per panicle**. Selection frequency across 100 LLM-guided ZSW runs for the initial field trials, shown on a principal component (PC) plot derived from marker genotypes (PC1 vs PC2). Here, we compared four models: (A) CropSeek, (B) Gemma, (C) Mistral, and (D) Openchat. As in Fig. 5, point color denotes rice subpopulation, and point size is proportional to the number of times each genotype was selected (0–100). Filled circles indicate genotypes selected at least once; open circles indicate genotypes never selected.

**Supplementary Fig. 4. Genotype selection frequency in seven zero-shot warmstarting models for protein content**. As in Fig. 5, the figure shows the selection frequency across 100 LLM-guided ZSW runs for the initial field trials on a principal component (PC) plot derived from marker genotypes (PC1 vs PC2). We compared seven models: (A) Llama, (B) PLLaMa, (C) Random, (D) CropSeek, (E) Gemma, (F) Mistral, and (G) Openchat. Point color denotes rice subpopulation, and point size is proportional to the number of times each genotype was selected (0–100). Filled circles indicate genotypes selected at least once; open circles indicate genotypes never selected.

**Supplementary Fig. 5. Comparison of the Mistral-based surrogate model (SGM) and the conventional SGM when genomic information is available: improvement in the maximum/minimum phenotypic value among selected genotypes for (A) seed number per panicle and (B) protein content**. As in Fig. 4, the x-axis denotes the number of genotypes already evaluated in field trials, and the y-axis denotes the maximum/minimum phenotypic value among the genotypes evaluated so far. Here, we compared three strategies using genomic information: LLG (purple), which uses an LLM (Mistral) for both ZSW and SGM; LG (light green), which uses Mistral for ZSW and a Gaussian process for SGM; and LRG (black), which uses Mistral for ZSW and Random for SGM.

**Supplementary Fig. 6. Comparison of the Llama-based surrogate model (SGM) and the conventional SGM when genomic information is available: change in the surrogate model accuracy for (A) seed number per panicle and (B) protein content**. As in Supplementary Fig. 2, the figure plots the change in the surrogate model accuracy (y-axis) against the number of genotypes evaluated in field trials so far (x-axis). Abbreviations and the line colors corresponding to each strategy are the same as in Fig. 6.

**Supplementary Fig. 7. Comparison of LLMs for zero-shot warmstarting (ZSW): change in the surrogate model accuracy for the trait simulated under (A) Scenario 1, (B) Scenario 2, and (C) Scenario 3**. As in Supplementary Fig. 2, the figure plots the change in the surrogate model accuracy (y-axis) against the number of genotypes evaluated in field trials so far (x-axis). Model abbreviations and the line colors corresponding to each model are the same as in Fig. 4.

**Supplementary Fig. 8. Genotype selection frequency in seven zero-shot warmstarting models for the trait simulated under Scenario 1**. As in Fig. 5, the figure shows the selection frequency across 100 LLM-guided ZSW runs for the initial field trials on a principal component (PC) plot derived from marker genotypes (PC1 vs PC2). We compared seven models: (A) Llama, (B) PLLaMa, (C) Random, (D) CropSeek, (E) Gemma, (F) Mistral, and (G) Openchat. Point color denotes rice subpopulation, and point size is proportional to the number of times each genotype was selected (0–100). Filled circles indicate genotypes selected at least once; open circles indicate genotypes never selected.

**Supplementary Fig. 9. Genotype selection frequency in seven zero-shot warmstarting models for the trait simulated under Scenario 2**. As in Fig. 5, the figure shows the selection frequency across 100 LLM-guided ZSW runs for the initial field trials on a principal component (PC) plot derived from marker genotypes (PC1 vs PC2). We compared seven models: (A) Llama, (B) PLLaMa, (C) Random, (D) CropSeek, (E) Gemma, (F) Mistral, and (G) Openchat. Point color denotes rice subpopulation, and point size is proportional to the number of times each genotype was selected (0–100). Filled circles indicate genotypes selected at least once; open circles indicate genotypes never selected.

**Supplementary Fig. 10. Genotype selection frequency in seven zero-shot warmstarting models for the trait simulated under Scenario 3**. As in Fig. 5, the figure shows the selection frequency across 100 LLM-guided ZSW runs for the initial field trials on a principal component (PC) plot derived from marker genotypes (PC1 vs PC2). We compared seven models: (A) Llama, (B) PLLaMa, (C) Random, CropSeek, (E) Gemma, (F) Mistral, and (G) Openchat. Point color denotes rice subpopulation, and point size is proportional to the number of times each genotype was selected (0–100). Filled circles indicate genotypes selected at least once; open circles indicate genotypes never selected.

**Table 1 Strategies compared in this study**.

**Supplementary Table 1. Comparison of zero-shot warmstarting models by the number of genotypes evaluated in field trials needed to identify the best genotype for seed number per panicle**.

**Supplementary Table 2. Comparison of zero-shot warmstarting models by the number of genotypes evaluated in field trials needed to identify the best genotype for protein content**.

**Supplementary Table 3. Comparison of the Llama-based surrogate model (SGM) and the conventional SGM with genomic information by the number of genotypes evaluated in field trials needed to identify the best genotype for seed number per panicle**.

**Supplementary Table 4. Comparison of the Llama-based surrogate model (SGM) and the conventional SGM with genomic information by the number of genotypes evaluated in field trials needed to identify the best genotype for protein content**.

**Supplementary Table 5. Comparison of the Mistral-based surrogate model (SGM) and the conventional SGM with genomic information by the number of genotypes evaluated in field trials needed to identify the best genotype for seed number per panicle**.

**Supplementary Table 6. Comparison of the Mistral-based surrogate model (SGM) and the conventional SGM with genomic information by the number of genotypes evaluated in field trials needed to identify the best genotype for protein content**.

**Supplementary Table 7. Comparison of the Llama-based surrogate model (SGM) and the conventional SGM without genomic information by the number of genotypes evaluated in field trials needed to identify the best genotype for seed number per panicle**.

**Supplementary Table 8. Comparison of the Llama-based surrogate model (SGM) and the conventional SGM without genomic information by the number of genotypes evaluated in field trials needed to identify the best genotype for protein content**.

## Declarations

### Ethics approval and consent to participate

Not applicable.

### Consent for publication

Not applicable.

### Availability of data and materials

All the scripts used in this study, including the LLM-guided Bayesian optimization, are available from the “KosukeHamazaki/RLLMBO” repository on GitHub, https://github.com/KosukeHamazaki/RLLMBO. All datasets, including the results, will be shared in the repository on Zenodo, https://doi.org/10.5281/zenodo.20824616.

### Authors’ contributions

KH and KT conceived the study of incorporating large language models into Bayesian optimization for agricultural problems. KH determined the detailed problem settings, preprocessed the open dataset, simulated the data, implemented the LLM-guided and conventional BOs, visualized the results, and wrote the draft. KT checked and revised the draft. All authors read and approved the final manuscript.

## Acknowledgements

The computations in this work were performed in the supercomputer centers at RAIDEN. We would like to thank Dr. Masato Sumita for his assistance in setting up the computational environment, such as RAIDEN.

## References

[1] Cooper, M., Messina, C.D.: Breeding crops for drought-affected environments and improved climate resilience. Plant Cell 35(1), 162–186 (2023)

[2] Mittler, R., Karlova, R., Bassham, D.C., Lawson, T.: Crops under stress: can we mitigate the impacts of climate change on agriculture and launch the ‘resilience revolution’? Philos. Trans. R. Soc. Lond. B Biol. Sci. 380(1927), 20240228 (2025)

[3] McCouch, S., Baute, G.J., Bradeen, J., Bramel, P., Bretting, P.K., Buckler, E., Burke, J.M., Charest, D., Cloutier, S., Cole, G., Dempewolf, H., Dingkuhn, M., Feuillet, C., Gepts, P., Grattapaglia, D., Guarino, L., Jackson, S., Knapp, S., Langridge, P., Lawton-Rauh, A., Lijua, Q., Lusty, C., Michael, T., Myles, S., Naito, K., Nelson, R.L., Pontarollo, R., Richards, C.M., Rieseberg, L., Ross-Ibarra, J., Rounsley, S., Hamilton, R.S., Schurr, U., Stein, N., Tomooka, N., Knaap, E., Tassel, D., Toll, J., Valls, J., Varshney, R.K., Ward, J., Waugh, R., Wenzl, P., Zamir, D.: Agriculture: Feeding the future: Agriculture. Nature 499(7456), 23–24 (2013)

[4] Dempewolf, H., Eastwood, R.J., Guarino, L., Khoury, C.K., Müller, J.V., Toll, J.: Adapting agriculture to climate change: A global initiative to collect, conserve, and use crop wild relatives. Agroecol. Sustain. Food Syst. 38(4), 369–377 (2014)

[5] Khoury, C.K., Brush, S., Costich, D.E., Curry, H.A., Haan, S., Engels, J.M.M., Guarino, L., Hoban, S., Mercer, K.L., Miller, A.J., Nabhan, G.P., Perales, H.R., Richards, C., Riggins, C., Thormann, I.: Crop genetic erosion: understanding and responding to loss of crop diversity. New Phytol. 233(1), 84–118 (2022)

[6] Salgotra, R.K., Chauhan, B.S.: Genetic diversity, conservation, and utilization of plant genetic resources. Genes (Basel) 14(1), 174 (2023)

[7] 3,000 rice genomes project: The 3,000 rice genomes project. Gigascience 3, 7 (2014)

[8] Wang, W., Mauleon, R., Hu, Z., Chebotarov, D., Tai, S., Wu, Z., Li, M., Zheng, T., Fuentes, R.R., Zhang, F., Mansueto, L., Copetti, D., Sanciangco, M., Palis, K.C., Xu, J., Sun, C., Fu, B., Zhang, H., Gao, Y., Zhao, X., Shen, F., Cui, X., Yu, H., Li, Z., Chen, M., Detras, J., Zhou, Y., Zhang, X., Zhao, Y., Kudrna, D., Wang, C., Li, R., Jia, B., Lu, J., He, X., Dong, Z., Xu, J., Li, Y., Wang, M., Shi, J., Li, J., Zhang, D., Lee, S., Hu, W., Poliakov, A., Dubchak, I., Ulat, V.J., Borja, F.N., Mendoza, J.R., Ali, J., Li, J., Gao, Q., Niu, Y., Yue, Z., Naredo, M.E.B., Talag, J., Wang, X., Li, J., Fang, X., Yin, Y., Glaszmann, J.-C., Zhang, J., Li, J., Hamilton, R.S., Wing, R.A., Ruan, J., Zhang, G., Wei, C., Alexandrov, N., McNally, K.L., Li, Z., Leung, H.: Genomic variation in 3,010 diverse accessions of asian cultivated rice. Nature 557(7703), 43–49 (2018)

[9] Grant, D., Nelson, R.T., Cannon, S.B., Shoemaker, R.C.: SoyBase, the USDA-ARS soybean genetics and genomics database. Nucleic Acids Res. 38(Database issue), 843–6 (2010)

[10] Brown, A.V., Conners, S.I., Huang, W., Wilkey, A.P., Grant, D., Weeks, N.T., Cannon, S.B., Graham, M.A., Nelson, R.T.: A new decade and new data at SoyBase, the USDA-ARS soybean genetics and genomics database. Nucleic Acids Res. 49(D1), 1496–1501 (2021)

[11] Nguyen, G.N., Norton, S.L.: Genebank phenomics: A strategic approach to enhance value and utilization of crop germplasm. Plants 9(7), 817 (2020)

[12] Khazaei, H., Street, K., Bari, A., Mackay, M., Stoddard, F.L.: The FIGS (focused identification of germplasm strategy) approach identifies traits related to drought adaptation in vicia faba genetic resources. PLoS One 8(5), 63107 (2013)

[13] Tanaka, R., Iwata, H.: Bayesian optimization for genomic selection: a method for discovering the best genotype among a large number of candidates. Theor. Appl. Genet. 131(1), 93–105 (2018)

[14] Tsai, S.-F., Shen, C.-C., Liao, C.-T.: Bayesian optimization approaches for identifying the best genotype from a candidate population. J. Agric. Biol. Environ. Stat. 26(4), 519–537 (2021)

[15] Brochu, E., Cora, V.M., Freitas, N.: A tutorial on bayesian optimization of expensive cost functions, with application to active user modeling and hierarchical reinforcement learning. arXiv [cs.LG] (2010) [cs.LG]

[16] Shahriari, B., Swersky, K., Wang, Z., Adams, R.P., Freitas, N.: Taking the human out of the loop: A review of bayesian optimization. Proc. IEEE Inst. Electr. Electron. Eng. 104(1), 148–175 (2016)

[17] Hamazaki, K., Iwata, H.: Bayesian optimization of multivariate genomic prediction models based on secondary traits for improved accuracy gains and phenotyping costs. Theor. Appl. Genet. 135(1), 35–50 (2022)

[18] Diot, J., Iwata, H.: Bayesian optimisation for breeding schemes. Front. Plant Sci. 13, 1050198 (2022)

[19] Jannink, J.-L., Astudillo, R., Frazier, P.: Insight into a two-part plant breeding scheme through bayesian optimization of budget allocations. Crop Sci. (2023)

[20] Tu, H.-N., Liao, C.-T.: A modified bayesian optimization approach for determining a training set to identify the best genotypes from a candidate population in genomic selection. J. Agric. Biol. Environ. Stat. 31(1), 1–22 (2026)

[21] Vaswani, A., Shazeer, N., Parmar, N., Uszkoreit, J., Jones, L., Gomez, A.N., Kaiser, L., Polosukhin, I.: Attention is all you need. arXiv [cs.CL] (2017) [cs.CL]

[22] Brown, T.B., Mann, B., Ryder, N., Subbiah, M., Kaplan, J., Dhariwal, P., Neelakantan, A., Shyam, P., Sastry, G., Askell, A., Agarwal, S., Herbert-Voss, A., Krueger, G., Henighan, T., Child, R., Ramesh, A., Ziegler, D.M., Wu, J., Winter, C., Hesse, C., Chen, M., Sigler, E., Litwin, M., Gray, S., Chess, B., Clark, J., Berner, C., McCandlish, S., Radford, A., Sutskever, I., Amodei, D.: Language models are few-shot learners. arXiv [cs.CL] (2020) [cs.CL]

[23] Devlin, J., Chang, M.-W., Lee, K., Toutanova, K.: BERT: Pre-training of deep bidirectional transformers for language understanding. In: Proceedings of the 2019 Conference of the North, pp. 4171–4186. Association for Computational Linguistics, Stroudsburg, PA, USA (2019)

[24] Lewis, P., Perez, E., Piktus, A., Petroni, F., Karpukhin, V., Goyal, N., Küttler, H., Lewis, M., Yih, W.-T., Rocktäschel, T., Riedel, S., Kiela, D.: Retrieval-augmented generation for knowledge-intensive NLP tasks. arXiv [cs.CL] (2020) [cs.CL]

[25] Zhu, H., Qin, S., Su, M., Lin, C., Li, A., Gao, J.: Harnessing large vision and language models in agriculture: a review. Front. Plant Sci. 16(1579355), 1579355 (2025)

[26] Yang, X., Gao, J., Xue, W., Alexandersson, E.: PLLaMa: An open-source large language model for plant science. arXiv [cs.CL] (2024) [cs.CL]

[27] Yang, B., Zhang, Y., Feng, L., Chen, Y., Zhang, J., Xu, X., Aierken, N., Li, Y., Chen, Y., Yang, G., He, Y., Huang, R., Li, S.: AgriGPT: A large language model ecosystem for agriculture. arXiv [cs.AI] (2025) [cs.AI]

[28] Silva, B., Nunes, L., Estevão, R., Aski, V., Chandra, R.: GPT-4 as an agronomist assistant? answering agriculture exams using large language models. arXiv [cs.AI] (2023) [cs.AI]

[29] Ying, J., Chen, Z., Wang, Z., Jiang, W., Wang, C., Yuan, Z., Su, H., Kong, H., Yang, F., Dong, N.: SeedBench: A multi-task benchmark for evaluating large language models in seed science. arXiv [cs.CL] (2025) [cs.CL]

[30] Zhang, R., Wang, Y., Yang, W., Wen, J., Liu, W., Zhi, S., Li, G., Chai, N., Huang, J., Xie, Y., Xie, X., Chen, L., Gu, M., Liu, Y.-G., Zhu, Q.: PlantGPT: An arabidopsis-based intelligent agent that answers questions about plant functional genomics. Adv. Sci. (Weinh.) 12(30), 03926 (2025)

[31] Yoosefzadeh-Najafabadi, M.: From text to traits: exploring the role of large language models in plant breeding. Front. Plant Sci. 16, 1583344 (2025)

[32] Yang, C., Wang, X., Lu, Y., Liu, H., Le, Q.V., Zhou, D., Chen, X.: Large language models as optimizers. arXiv [cs.LG] (2023) [cs.LG]

[33] Liu, S., Chen, C., Qu, X., Tang, K., Ong, Y.-S.: Large language models as evolutionary optimizers. arXiv [cs.NE] (2023) [cs.NE]

[34] Liu, T., Astorga, N., Seedat, N., Schaar, M.: Large language models to enhance bayesian optimization. arXiv [cs.LG] (2024) [cs.LG]

[35] Rychert, A., Spagnolo, G., Posashkov, E.: Reproducibility study of large language 29 model bayesian optimization. arXiv [cs.CL] (2025) [cs.CL]

[36] Yang, Z., Wang, D., Ge, L., Wang, B., Fu, T., Li, Y.: Reasoning BO: Enhancing bayesian optimization with long-context reasoning power of LLMs. arXiv [cs.AI] (2025) [cs.AI]

[37] Kristiadi, A., Strieth-Kalthoff, F., Skreta, M., Poupart, P., Aspuru-Guzik, A., Pleiss, G.: A sober look at LLMs for material discovery: Are they actually good for bayesian optimization over molecules? arXiv [cs.LG] (2024) [cs.LG]

[38] Yin, Y., Wang, Y., Xu, B., Li, P.: ADO-LLM: Analog design bayesian optimization with in-context learning of large language models. arXiv [cs.LG] (2024) [cs.LG]

[39] Yuan, X., Chen, Z., Zhang, J., Xiong, H., Ye, N., Li, Y., Gu, Q.: Unleashing LLMs in bayesian optimization: Preference-guided framework for scientific discovery. arXiv [cs.AI] (2026) [cs.AI]

[40] R Core Team: R: A Language and Environment for Statistical Computing. R Foundation for Statistical Computing, Vienna, Austria (2025). R Foundation for Statistical Computing. https://www.R-project.org/

[41] Wickham, H.: Ggplot2: Elegant Graphics for Data Analysis, 2nd edn. Use R! Springer, Cham, Switzerland (2016)

[42] Rasmussen, C.E., Williams, C.K.I.: Gaussian Processes for Machine Learning. Adaptive Computation and Machine Learning series. MIT Press, London, England (2005)

[43] Gardner, J.R., Pleiss, G., Bindel, D., Weinberger, K.Q., Wilson, A.G.: GPyTorch: Blackbox matrix-matrix gaussian process inference with GPU acceleration. arXiv [cs.LG] (2018) [cs.LG]

[44] Srinivas, N., Krause, A., Kakade, S.M., Seeger, M.: Gaussian process optimization in the bandit setting: No regret and experimental design. arXiv [cs.LG] (2009) [cs.LG]

[45] Zhao, K., Tung, C.-W., Eizenga, G.C., Wright, M.H., Ali, M.L., Price, A.H., Norton, G.J., Islam, M.R., Reynolds, A., Mezey, J., McClung, A.M., Bustamante, C.D., McCouch, S.R.: Genome-wide association mapping reveals a rich genetic architecture of complex traits in oryza sativa. Nat. Commun. 2, 467 (2011)

[46] Purcell, S., Neale, B., Todd-Brown, K., Thomas, L., Ferreira, M.A.R., Bender, D., Maller, J., Sklar, P., Bakker, P.I.W., Daly, M.J., Sham, P.C.: PLINK: a tool set for whole-genome association and population-based linkage analyses. Am. J. Hum. Genet. 81(3), 559–575 (2007)

[47] Chang, C.C., Chow, C.C., Tellier, L.C., Vattikuti, S., Purcell, S.M., Lee, J.J.: Second-generation PLINK: rising to the challenge of larger and richer datasets. Gigascience 4, 7 (2015)

[48] Danecek, P., Auton, A., Abecasis, G., Albers, C.A., Banks, E., DePristo, M.A., Handsaker, R.E., Lunter, G., Marth, G.T., Sherry, S.T., McVean, G., Durbin, R., 1000 Genomes Project Analysis Group: The variant call format and VCFtools. Bioinformatics 27(15), 2156–2158 (2011)

[49] Browning, S.R., Browning, B.L.: Rapid and accurate haplotype phasing and missing-data inference for whole-genome association studies by use of localized haplotype clustering. Am. J. Hum. Genet. 81(5), 1084–1097 (2007)

[50] Browning, B.L., Browning, S.R.: Genotype imputation with millions of reference samples. Am. J. Hum. Genet. 98(1), 116–126 (2016)

[51] Persadh, D. Darshani .R: CropSeek-LLM: Agricultural Domain Language Model. Hugging Face (2025). 10.57967/hf/5849

[52] Gemma Team: Thomas Mesnard, Hardin, C., Dadashi, R., Bhupatiraju, S., Pathak, S., Sifre, L., Rivière, M., Kale, M.S., Love, J., Tafti, P., Hussenot, L., Sessa, P.G., Chowdhery, A., Roberts, A., Barua, A., Botev, A., Castro-Ros, A., Slone, A., Héliou, A., Tacchetti, A., Bulanova, A., Paterson, A., Tsai, B., Shahriari, B., Le Lan, C., Choquette-Choo, C.A., Crepy, C., Cer, D., Ippolito, D., Reid, D., Buchatskaya, E., Ni, E., Noland, E., Yan, G., Tucker, G., Muraru, G.-C., Rozhdestvenskiy, G., Michalewski, H., Tenney, I., Grishchenko, I., Austin, J., Keeling, J., Labanowski, J., Lespiau, J.-B., Stanway, J., Brennan, J., Chen, J., Ferret, J., Chiu, J., Mao-Jones, J., Lee, K., Yu, K., Millican, K., Sjoesund, L.L., Lee, L., Dixon, L., Reid, M., Mikula, M., Wirth, M., Sharman, M., Chinaev, N., Thain, N., Bachem, O., Chang, O., Wahltinez, O., Bailey, P., Michel, P., Yotov, P., Chaabouni, R., Comanescu, R., Jana, R., Anil, R., McIlroy, R., Liu, R., Mullins, R., Smith, S.L., Borgeaud, S., Girgin, S., Douglas, S., Pandya, S., Shakeri, S., De, S., Klimenko, T., Hennigan, T., Feinberg, V., Stokowiec, W., Chen, Y.-H., Ahmed, Z., Gong, Z., Warkentin, T., Peran, L., Giang, M., Farabet, C., Vinyals, O., Dean, J., Kavukcuoglu, K., Hassabis, D., Ghahramani, Z., Eck, D., Barral, J., Pereira, F., Collins, E., Joulin, A., Fiedel, N., Senter, E., Andreev, A., Kenealy, K.: Gemma: Open models based on gemini research and technology. arXiv [cs.CL] (2024) [cs.CL]

[53] Touvron, H., Lavril, T., Izacard, G., Martinet, X., Lachaux, M.-A., Lacroix, T., Rozière, B., Goyal, N., Hambro, E., Azhar, F., Rodriguez, A., Joulin, A., Grave, E., Lample, G.: LLaMA: Open and efficient foundation language models. arXiv [cs.CL] (2023) [cs.CL]

[54] Jiang, A.Q., Sablayrolles, A., Mensch, A., Bamford, C., Chaplot, D.S., Casas, D., Bressand, F., Lengyel, G., Lample, G., Saulnier, L., Lavaud, L.R., Lachaux, M.-A., Stock, P., Le Scao, T., Lavril, T., Wang, T., Lacroix, T., El Sayed, W.: Mistral 7B. arXiv [cs.CL] (2023) [cs.CL]

[55] Wang, G., Cheng, S., Zhan, X., Li, X., Song, S., Liu, Y.: OpenChat: Advancing open-source language models with mixed-quality data. arXiv [cs.CL] (2023) [cs.CL]

[56] Bi, D.-A.X., Chen, D., Chen, G., Chen, S., Dai, D., Deng, C., Ding, H., Dong, K., Du, Q., Fu, Z., Gao, H., Gao, K., Gao, W., Ge, R., Guan, K., Guo, D., Guo, J., Hao, G., Hao, Z., He, Y., Hu, W., Huang, P., Li, E., Li, G., Li, J., Li, Y., Li, Y.K., Liang, W., Lin, F., Liu, A.X., Liu, B., Liu, W., Liu, X., Liu, X., Liu, Y., Lu, H., Lu, S., Luo, F., Ma, S., Nie, X., Pei, T., Piao, Y., Qiu, J., Qu, H., Ren, T., Ren, Z., Ruan, C., Sha, Z., Shao, Z., Song, J., Su, X., Sun, J., Sun, Y., Tang, M., Wang, B., Wang, P., Wang, S., Wang, Y., Wang, Y., Wu, T., Wu, Y., Xie, X., Xie, Z., Xie, Z., Xiong, Y., Xu, H., Xu, R.X., Xu, Y., Yang, D., You, Y., Yu, S., Yu, X., Zhang, B., Zhang, H., Zhang, L., Zhang, L., Zhang, M., Zhang, M., Zhang, W., Zhang, Y., Zhao, C., Zhao, Y., Zhou, S., Zhou, S., Zhu, Q., Zou, Y.: DeepSeek LLM: Scaling open-source language models with longtermism. arXiv [cs.CL] (2024) [cs.CL]

[57] Paszke, A., Gross, S., Massa, F., Lerer, A., Bradbury, J., Chanan, G., Killeen, T., Lin, Z., Gimelshein, N., Antiga, L., Desmaison, A., Köpf, A., Yang, E., DeVito, Z., Raison, M., Tejani, A., Chilamkurthy, S., Steiner, B., Fang, L., Bai, J., Chintala, S.: PyTorch: An imperative style, high-performance deep learning library. Adv. Neural Inf. Process. Syst. abs/1912.01703 (2019)

[58] Wolf, T., Debut, L., Sanh, V., Chaumond, J., Delangue, C., Moi, A., Cistac, P., Rault, T., Louf, R., Funtowicz, M., Davison, J., Shleifer, S., Platen, P., Ma, C., Jernite, Y., Plu, J., Xu, C., Le Scao, T., Gugger, S., Drame, M., Lhoest, Q., Rush, A.M.: HuggingFace’s transformers: State-of-the-art natural language processing. arXiv [cs.CL] (2019) [cs.CL]

[59] Reimers, N., Gurevych, I.: Sentence-BERT: Sentence embeddings using siamese BERT-networks. arXiv [cs.CL] (2019) [cs.CL]

[60] Pedregosa, F., Varoquaux, G., Gramfort, A., Michel, V., Thirion, B., Grisel, O., Blondel, M., Müller, A., Nothman, J., Louppe, G., Prettenhofer, P., Weiss, R., Dubourg, V., Vanderplas, J., Passos, A., Cournapeau, D., Brucher, M., Perrot, M., Duchesnay, E.: Scikit-learn: Machine learning in python. arXiv [cs.LG] (2012) [cs.LG]

[61] Ouyang, L., Wu, J., Jiang, X., Almeida, D., Wainwright, C.L., Mishkin, P., Zhang, C., Agarwal, S., Slama, K., Ray, A., Schulman, J., Hilton, J., Kelton, F., Miller, L., Simens, M., Askell, A., Welinder, P., Christiano, P., Leike, J., Lowe, R.: Training language models to follow instructions with human feedback. arXiv [cs.CL] (2022) [cs.CL]

[62] Luo, Y., Yang, Z., Meng, F., Li, Y., Zhou, J., Zhang, Y.: An empirical study 32 of catastrophic forgetting in large language models during continual fine-tuning. arXiv [cs.CL] (2023) [cs.CL]

[63] Zhang, S., Dong, L., Li, X., Zhang, S., Sun, X., Wang, S., Li, J., Hu, R., Zhang, T., Wu, F., Wang, G.: Instruction tuning for large language models: A survey. arXiv [cs.CL] (2023) [cs.CL]

[64] Consens, M.E., Dufault, C., Wainberg, M., Forster, D., Karimzadeh, M., Goodarzi, H., Theis, F.J., Moses, A., Wang, B.: To transformers and beyond: Large language models for the genome. arXiv [q-bio.GN] (2023) [q-bio.GN]

[65] Benegas, G., Ye, C., Albors, C., Li, J.C., Song, Y.S.: Genomic language models: opportunities and challenges. Trends Genet. 41(4), 286–302 (2025)

[66] Alagarasan, G., Li, H., Xu, Y., Crossa, J., Varshney, R.K.: Genomic language model-based genomic prediction in plant breeding. Trends Plant Sci. (2026)

[67] Adachi, M., Planden, B., Howey, D.A., Osborne, M.A., Orbell, S., Ares, N., Muandet, K., Chau, S.L.: Looping in the human collaborative and explainable bayesian optimization. arXiv [cs.LG] (2023) [cs.LG]

[68] Chakraborty, T., Wirth, C., Seifert, C.: Explainable bayesian optimization. arXiv [cs.LG] (2024) [cs.LG]

