## Supplementary figures and images for "Knowledge-guided Bayesian optimization using pre-trained LLMs speeds up the identification of superior genotypes from germplasm collection"

### Supplementary Figure 1

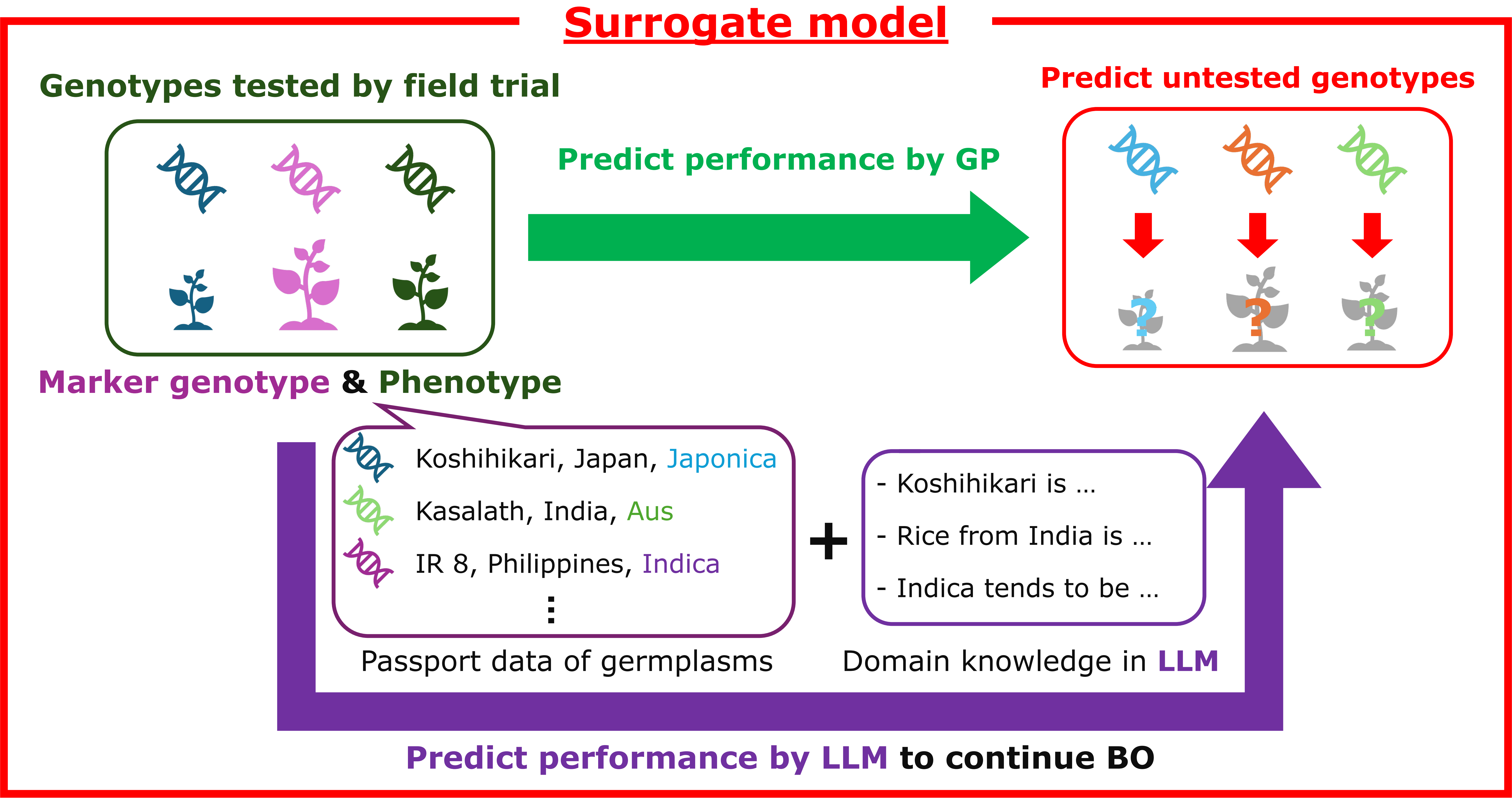

### Supplementary Figure 2

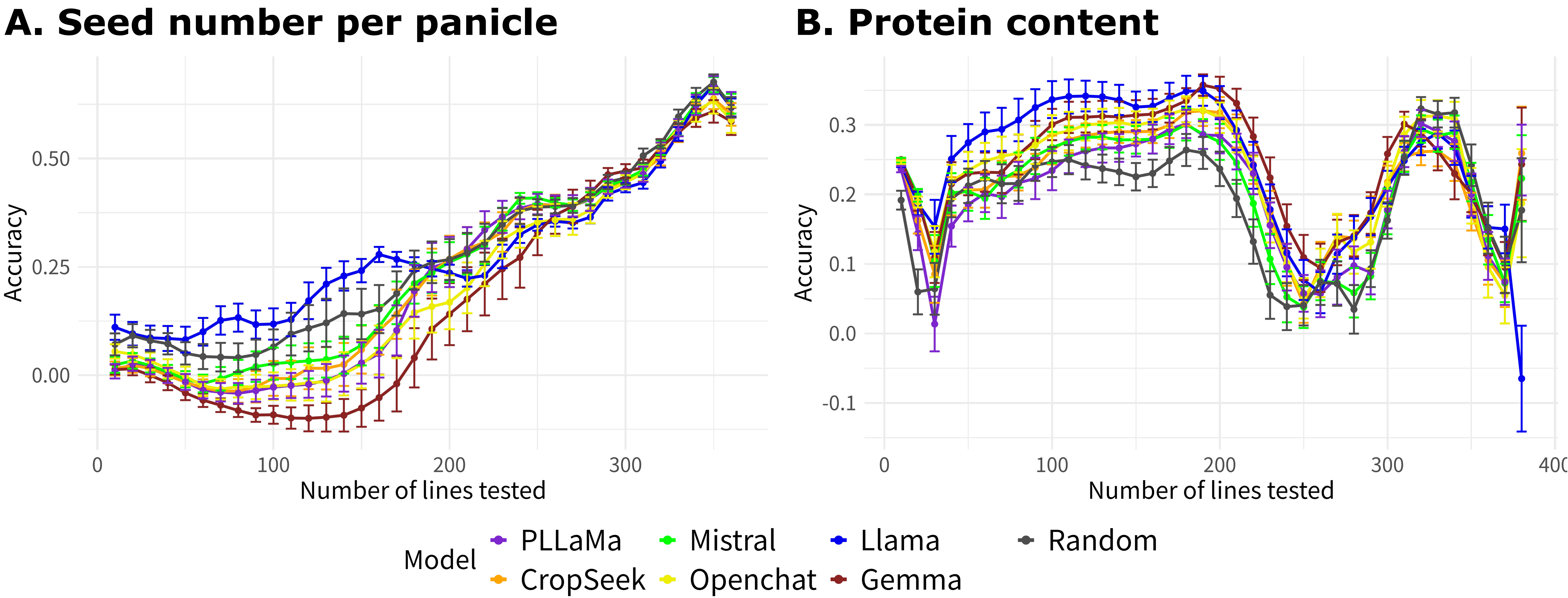

### Supplementary Figure 3

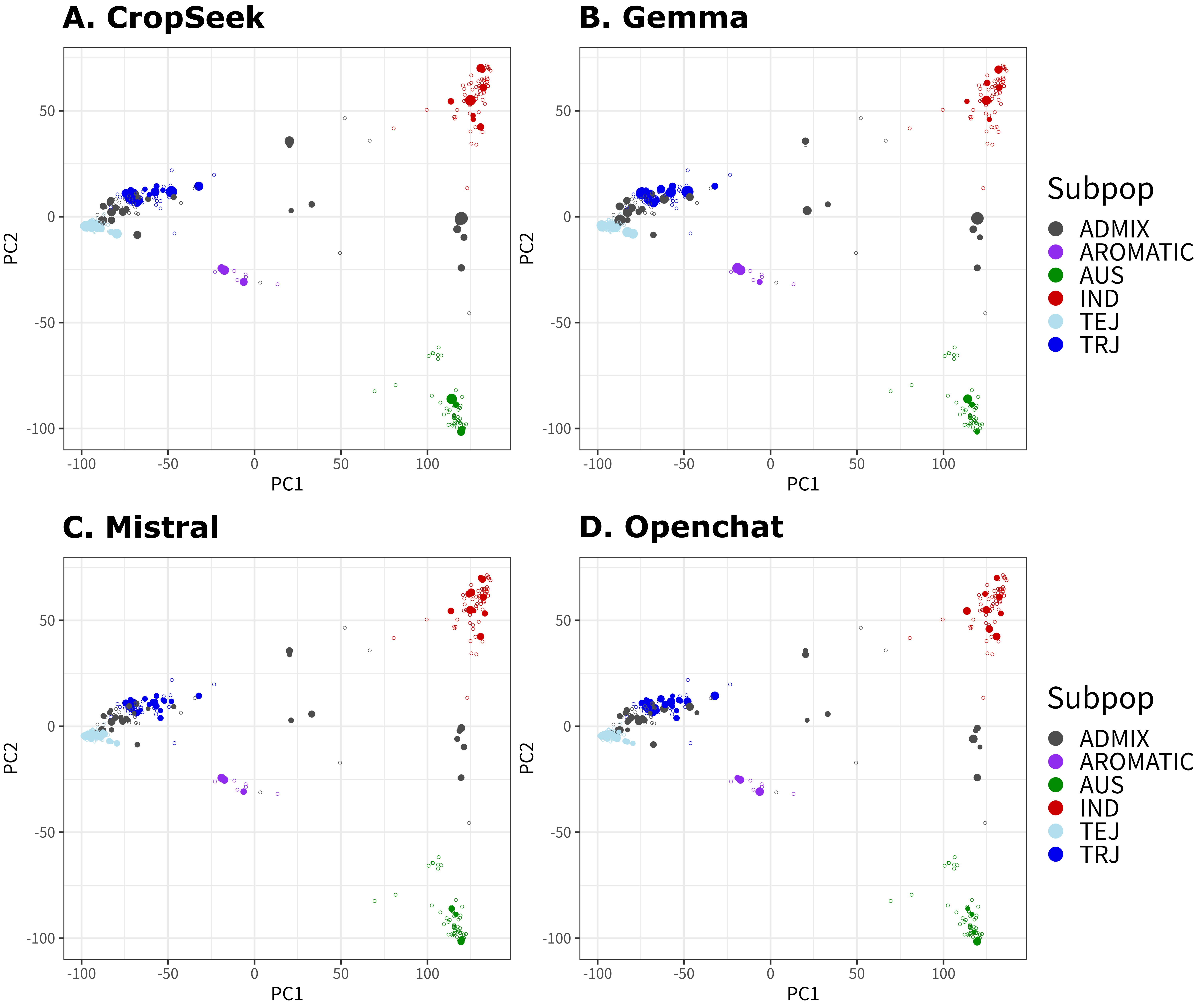

### Supplementary Figure 4

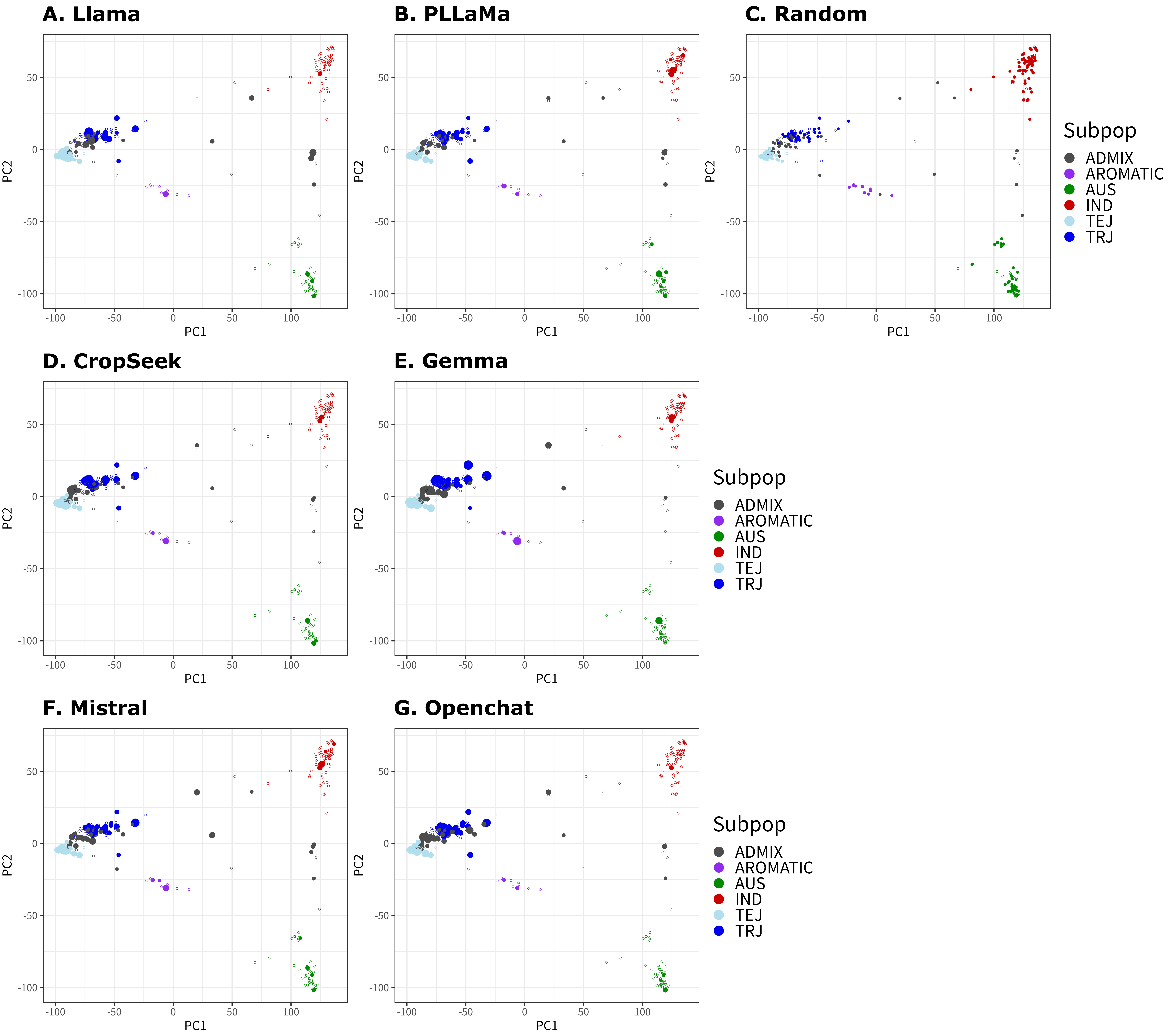

### Supplementary Figure 5

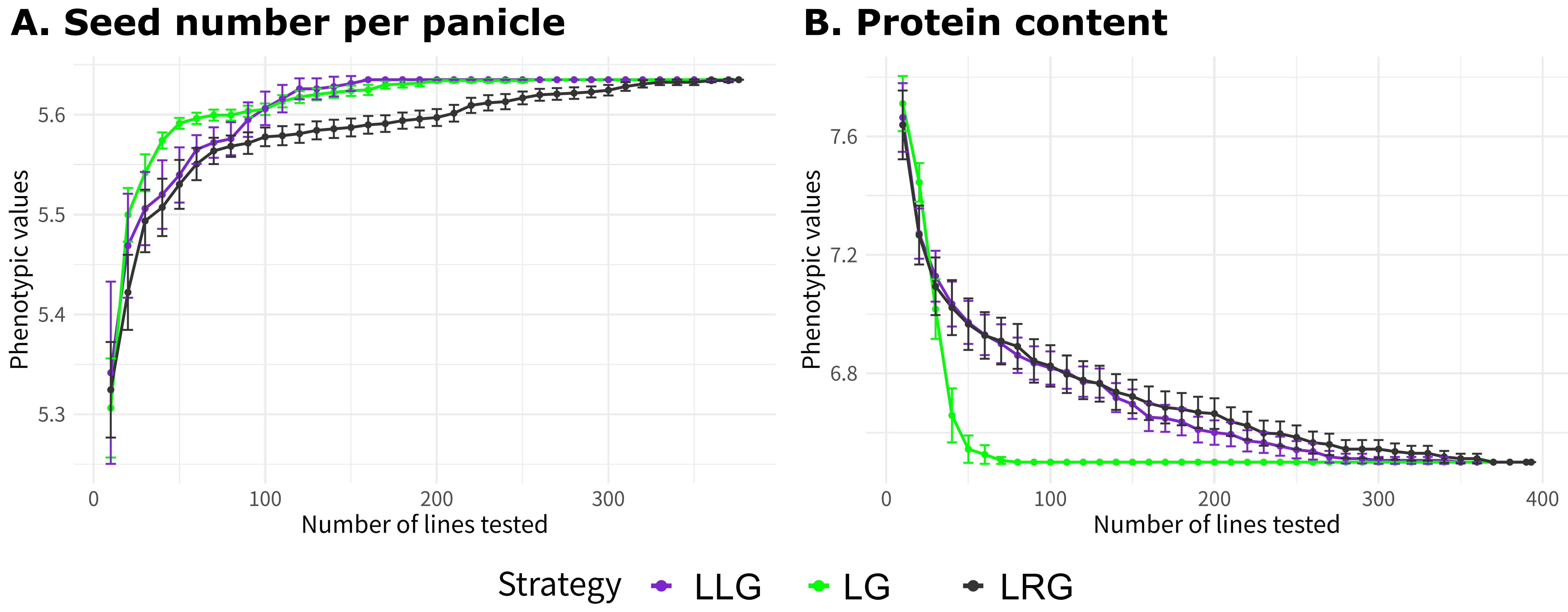

### Supplementary Figure 6

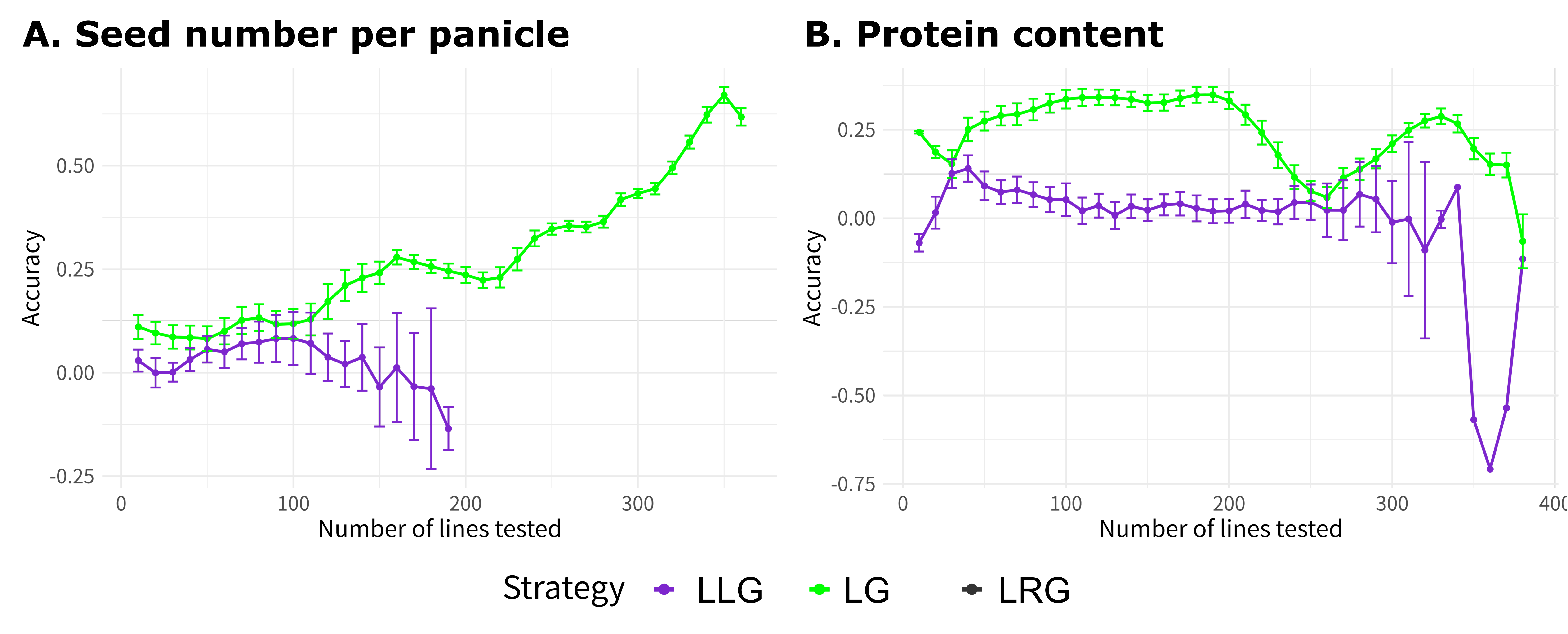

### Supplementary Figure 7

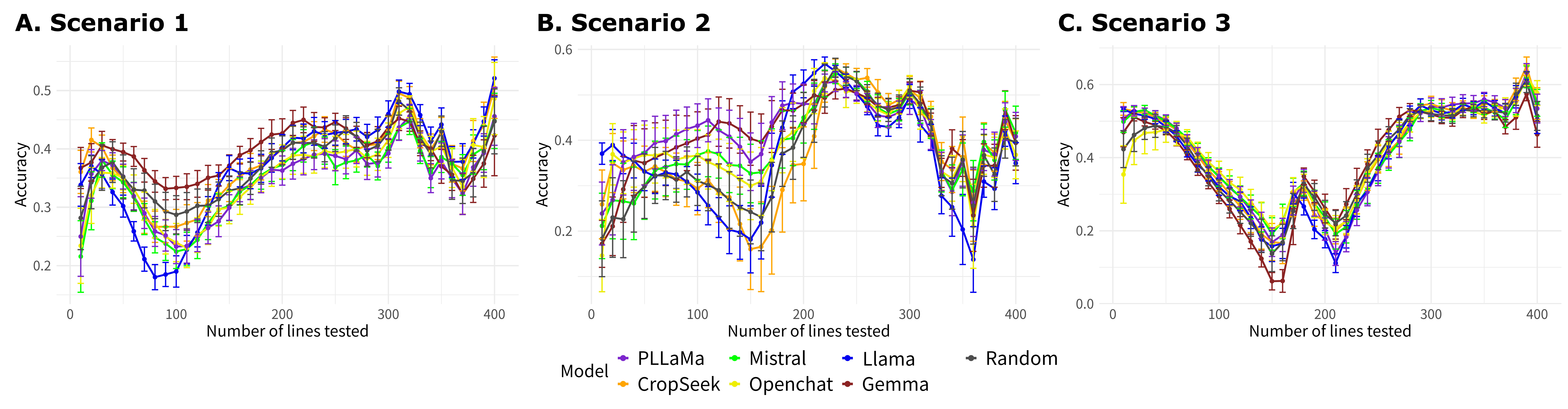

### Supplementary Figure 8

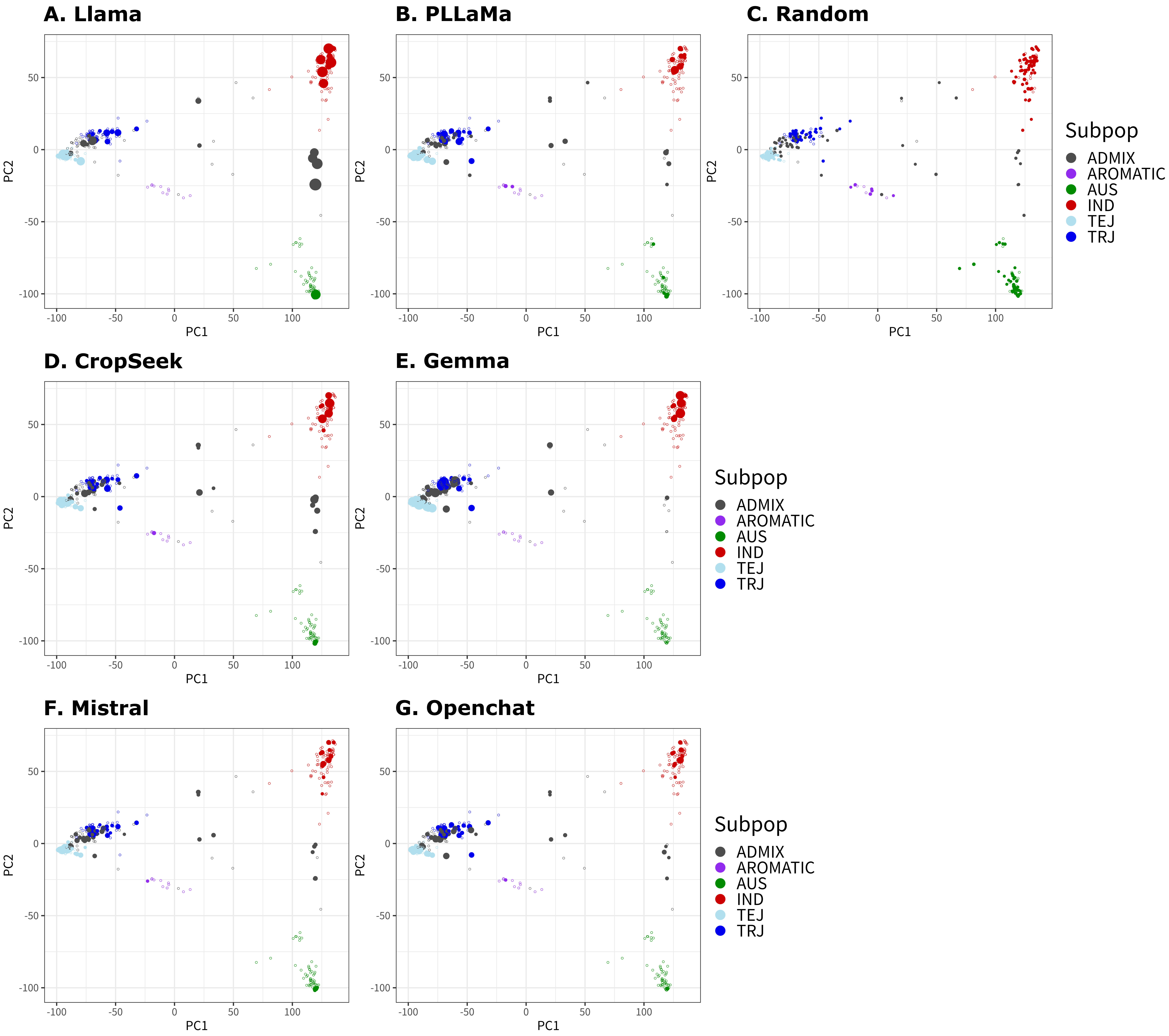

### Supplementary Figure 9

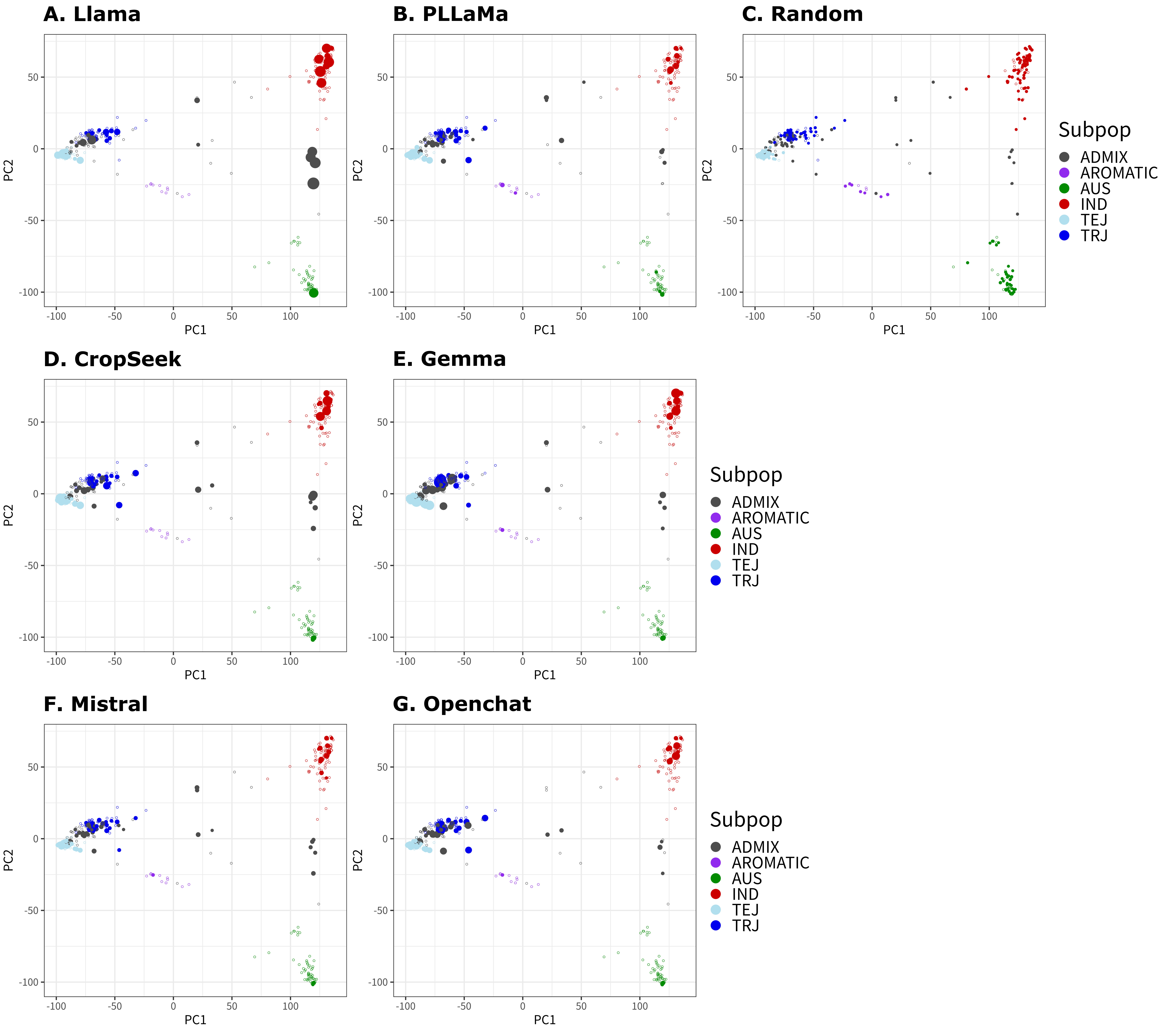

### Supplementary Figure 10

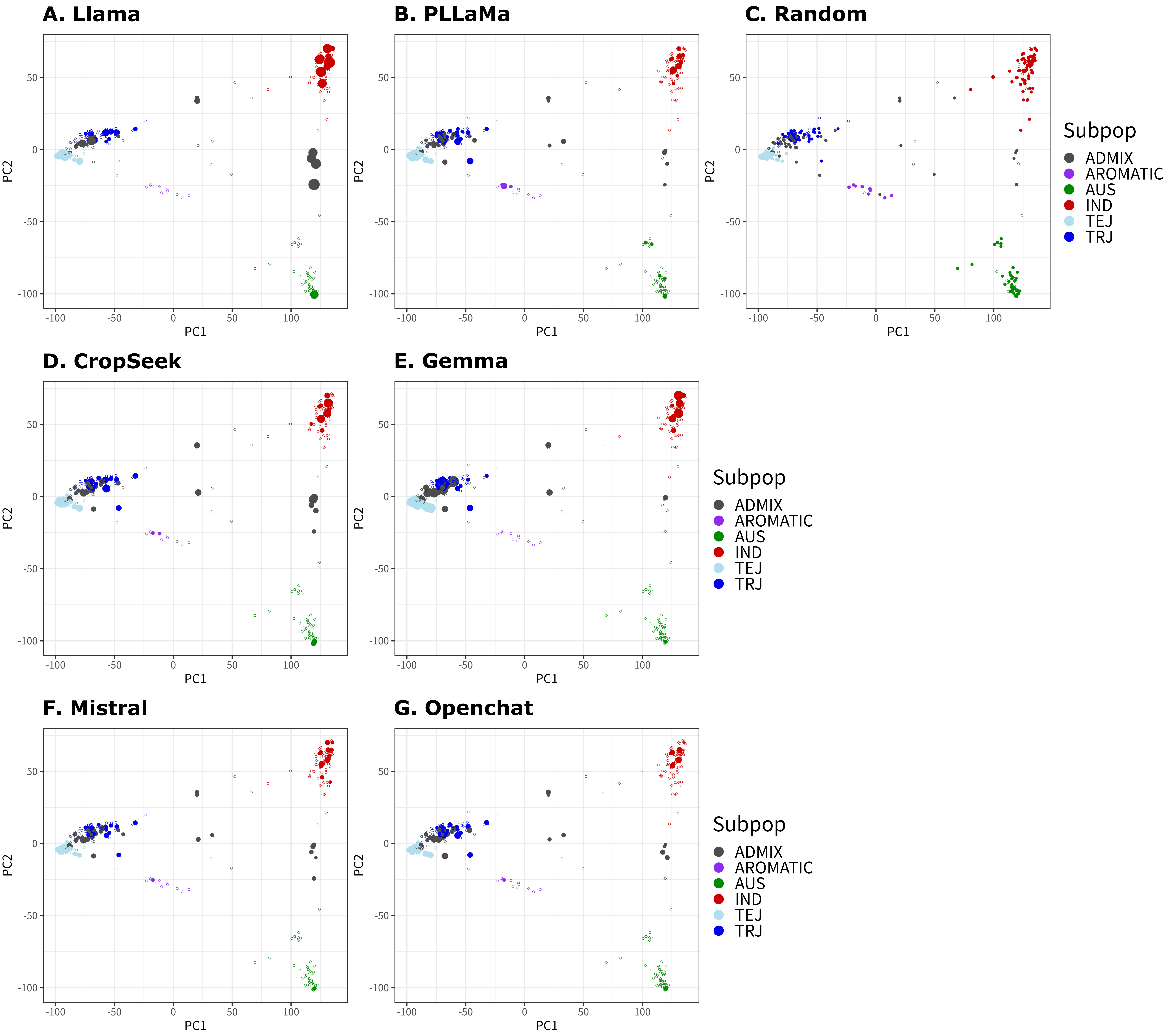
